# Pooled overexpression screening identifies PIPPI as a novel microprotein involved in the ER stress response

**DOI:** 10.1101/2024.12.08.627409

**Authors:** Lorenzo Lafranchi, Anna Spinner, Maximilian Hornisch, Dörte Schlesinger, Carmen Navarro Luzon, Linus Brinkenstråhle, Rui Shao, Ilaria Piazza, Simon J Elsässer

## Abstract

Microproteins encoded by short open reading frames (sORFs) of less than 100 codons have been predicted to constitute a substantial fraction of the eukaryotic proteome. However, relevance and roles of the majority of microproteins remain undefined because only a small fraction of these intriguing cellular players have been in-depth characterized so far. Here we use pooled overexpression screens with a library of 11’338 sORFs to overcome the challenge of elucidating which of the thousands of putative translated sORFs are biologically functional. As a proof-of-concept, we performed a phenotypic screen to identify sORFs protecting cells from treatment with the nucleotide analogue 6-thioguanine. With this approach, we identified two cytoprotective microproteins: altDDIT3 and PIPPI. PIPPI is encoded as part of the LC16a core duplicon/Morpheus gene cluster, a highly duplicated region of the human genome, which is undergoing rapid positive selection in primates. Our data show that PIPPI interacts with proteins of the endoplasmic reticulum, including protein disulfide isomerase ERp44. Besides providing mechanistic insights on a new microprotein, this study highlights the power of using pooled overexpression screens to identify functional microproteins.

## Introduction

Short open reading frames (sORFs) with the potential of encoding functional polypeptides have remained unnoticed for a long time due to their small size (i.e. less than 300 bp). Nevertheless, increasing evidence highlights that these microproteins, or micropeptides, are abundantly expressed in prokaryotic and eukaryotic cells. sORFs can be present in mRNA molecules, as an alternative to the main ORF, but also in other transcripts, such as long non-coding RNAs, microRNAs or antisense transcripts ^1,2^. Over the last years, a large amount of work has been invested in distinguishing between non-coding and translating sORFs. Ribosome profiling (Ribo-seq), mass spectrometry or bioinformatic assessment of evolutionary conservation are the main approaches used for identifying putative microproteins^1^. Finally, ongoing efforts are aiming to benchmark the ever-growing number of sORFs predicted to be translated using Ribo-seq and propose a high-confidence and standardized catalogue of human sORFs ^3^. Although thousands of microproteins have been predicted to exist in mammalian cells, only a small fraction of them is so far being thoroughly characterized. To mention a few prominent examples, microproteins have been shown to be involved in the maintenance of genome stability ^4–6^, gene expression ^7,8^, metabolism ^9,10^, and signaling ^11–14^. As it can be appreciated from this short list,microproteins display an intriguing variety of target tissues, subcellular compartments, and intra- or extracellular mechanisms of action ^15^. Ongoing research is highlighting how microproteins are not specifically evolved to solely serve in a subset of biological processes, but they appear to play an underappreciated role in cellular and organismal biology ^15,16^.

Cells are equipped with a plethora of evolutionary conserved signaling pathways that can be activated to counteract endogenous and exogenous stress events, thereby ensuring the maintenance of cellular homeostasis. In case of excessive stress, pro-death pathways represent a last resource used to remove severely damaged cells from the organism ^17^. Recent ribosome profiling studies highlighted that translation of a large number of sORFs, in particular upstream ORFs (uORFs), is upregulated in stress conditions ^18–21^. Despite some of these translational events could lack function at the protein level, we can expect that many functional stress-induced microproteins await to be discovered and characterized ^22^. Up to today, two of the major challenges in the field remain to elucidate which of the thousands of putative translated sORFs are biologically active and to functionally characterize them. Here we sought to identify bioactive microproteins with a pooled overexpression screen of sORFs encoded in the human genome. Introducing the sORF library into A375 cells, we seleted for increased resistance to the chemotherapeutic compound 6-thioguanine and identified two sORFs promoting cell proliferation and followed up with the in-depth characterization of one of them.

## Results

### Pooled overexpression screens to identify functional sORFs

Although thousands of microproteins have been predicted to exist in mammalian cells, it is highly challenging to define which of these putative microproteins have a function and to identify their biological role. To efficiently pinpoint functional microproteins, we set out to perform overexpression screens using a large library of sORFs encoding for putative microproteins. A comprehensive library was assembled as detailed in ^33^. An oligo pool of 11’338 sORFs coding for microproteins between 10 and 57 amino acids was gene-synthetized and cloned into a lentiviral vector (Figure 1A).

**Figure 1:**
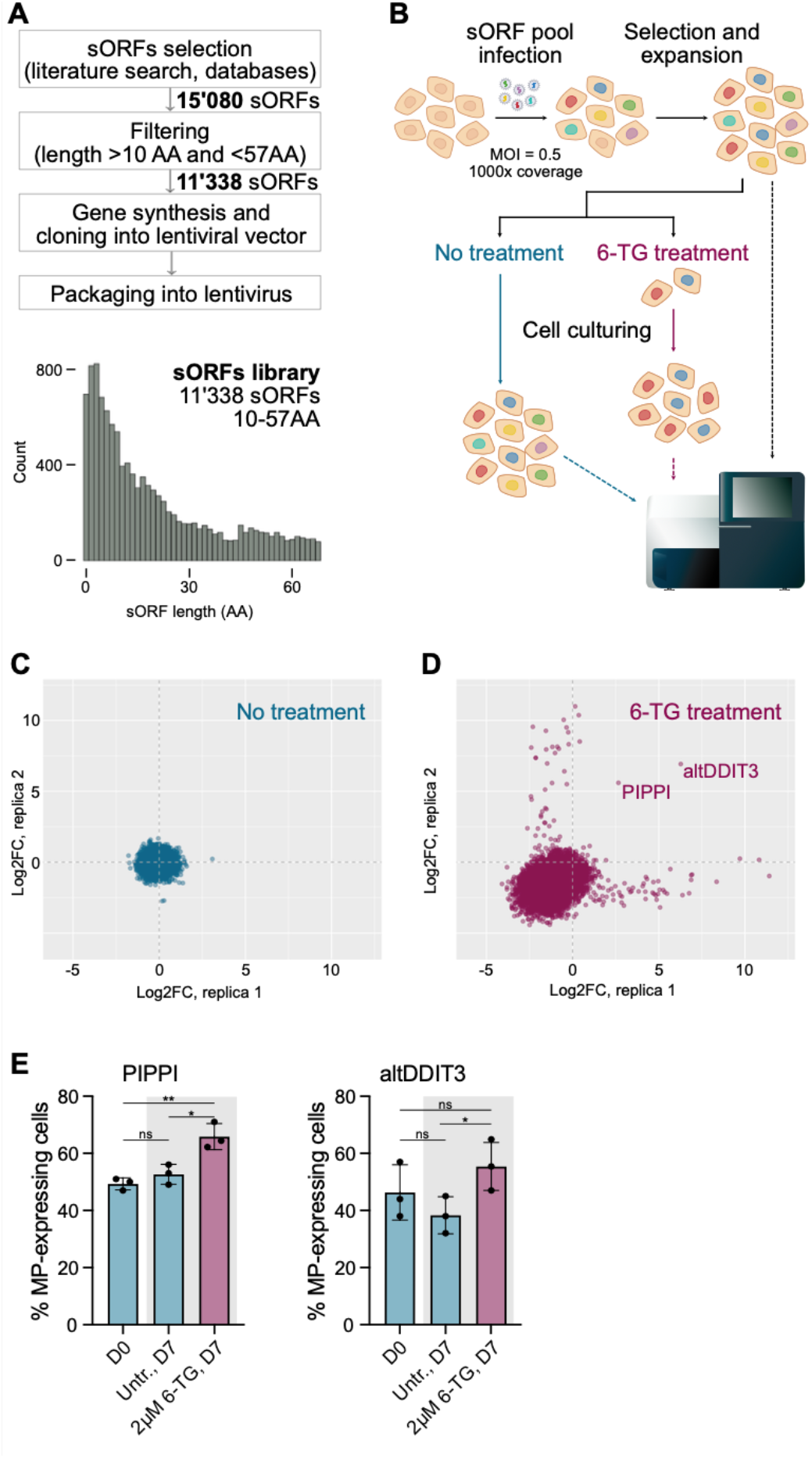
A pooled sORFs overexpression screen identifies sORFs conferring resistance to 6-thioguanine treatment. A) Schema depicting the rationale for the sORFs library design. B) Workflow of the pooled overexpression screens performed to identify sORFs modulating cell proliferation or resistance to 6-thioguanine (6-TG). Illustrations from bioicons.com were used as a basis for the graphics. C-D) Scatterplots showing the results of the cell proliferation screen (C) and the 6-TG screen (D). The log2 fold change (log2FC) in abundance of each sORF between the start and the end of the screen is plotted for the two biological replicates. E) Bar charts presenting the results of the growth competition assays performed to validate the results of the 6-TG screen. Where indicated, cells were treated for 24 hours with 2 µM 6-TG. Height of the bars represents the fraction of microprotein-expressing cells present in the total cell population. Values were averaged based on 3 independent biological replicates (black circles). p-values were calculated using unpaired Student’s t-test (ns = p > 0.05; * = p < 0.05; ** = p < 0.01).

Two screens were conducted in parallel in human A375 melanoma cells. In the first arm, cells were cultured untreated for three weeks to assess whether overexpression of any of the sORFs would promote cellular proliferation, whereas in the second arm of the screen cells were treated with the cytotoxic compound 6-thioguanine (6-TG). 6-TG is an analogue of the naturally occurring purine base guanine and it is mainly used in the clinic for the treatment of acute and chronic myelogenous leukemias. *In vivo,* 6-TG is converted to 6-Thioguanine nucleotides, which interfere with a variety of cellular processes involved in nucleic acid synthesis ^34^. For example, 6-TG nucleotides are incorporated in the genome of target cells during replication, and repair of the resulting lesions leads to cellular death. In this arm of the screen, cell cultures were continuously treated with 10 µM 6-TG until the surviving cells were able to repopulate the culturing flasks.

For the screen, over 10 Mio cells were transduced with the pool of sORFs at a low multiplicity of infection (MOI), ensuring that each sORF would be represented at least 1000 times in the cell population. After antibiotic selection to remove uninfected cells, a fraction of cells representing the starting population were collected for later extraction of genomic DNA. Thereafter, cells were cultured in absence or presence of 6-TG and, at the end of the screen, genomic DNA of the different populations was harvested. Finally, sORF cassettes were amplified by PCR and, after library preparation, the abundance of the individual sORFs in the different populations was quantified by Illumina sequencing (Figure 1B). By plotting the logarithmic fold-change (Log2 FC) in sORFs abundance between the start and the end populations of the cell proliferation screen, we could appreciate that overexpression of the different sORFs did not influence cell growth of normally proliferating cells (Figure 1C). Differently, in the 6-TG screen we could clearly see that a large fraction of sORFs had a negative Log2 FC value in both replicates of the screen indicating that the treatment killed the majority of cells (Figure 1D). Interestingly, two sORFs were on average 4-, respectively 7-fold enriched in both replicates upon drug treatment, suggesting that cells expressing these two microproteins are less sensitive toward 6-TG (Figure 1D). Beside these two hits, other sORFs were enriched in the final population, but only in one of the two replicates (Figure 1D). This was most likely occurring because of a proliferative advantage that few cells acquired thanks to random mutations, rather than the presence of the sORFs. The top hit corresponds to altDDIT3, which is an annotated microprotein encoded by a short open reading frame located upstream of the pro-apoptotic transcription factor CHOP/DDIT3 ^35–37^. Previous work on this uORF suggests that it can be translated to a functional microprotein that interacts with the downstream-encoded protein CHOP/DDIT3 ^35,38^. Nevertheless, it has also been shown that the uORF encoding for altDDIT3 principally acts *in cis*, either in a peptide-dependent or - independent manner, to downregulate translation of the canonical ORF ^39–42^. The second hit encoded a microprotein originally identified in a proteomics study ^43^. Matching sORF sequence with multiple possible non-canonical start codons were found 35 times in the human genome, and most of the matches were located as alternative ORFs on transcripts encoding the *nuclear pore complex interacting protein* (NPIP) family ^44,45^. We hence decided to dub this microprotein *PIPPI*.

To validate the results of the screen, we performed growth competition assays in which the proliferative capacity of a microprotein-overexpressing cell line was directly compared to that of its parental counterpart. In short, a microprotein-expressing cell line was mixed at a 1:1 ratio with a control cell line and the cell mixture was then either cultured untreated for one week or let recover from a 6-TG treatment for the same period. At the beginning and at the end of the assay, the ratio between GFP-positive and -negative cells was assessed by flow cytometry (Supplementary Figure 1A). Validating the screening results, expression of untagged PIPPI did not provide a proliferation advantage over GFP-only-expressing cells in untreated conditions but cells expressing the microprotein proliferated better than the parental cell line after treatment with 2 µM 6-TG (Figure 1E). Similarly, cells expressing altDDIT3 proliferated better than parental cells after a 6-TG treatment but not in untreated conditions (Figure 1E). The same observation could be made when performing the assay using a cell line expressing a GFP-tagged version of PIPPI and treating them with a higher dose of 6-TG (10 µM 6-TG, the same dose used for the screen). Interestingly, when the same strategy was applied to altDDIT3, we could observe that including the GFP tag inhibited the protective activity of altDDIT3 (Supplementary Figure 1B). Altogether, our data support previous reports indicating that altDDIT3 can be functional as a microprotein and suggest that it could have a role in inhibiting the pro-apoptotic effects of CHOP/altDDIT3, while PIPPI appeared to exert a similar effect through a yet unknown mechanism.

### PIPPI is a novel microprotein localizing to the ER

23 sORFs matching the PIPPI microprotein sequence were found on chr16 of the Ensemble 110 human genome annotation (Supplementary Table 2, Supplementary Information)^46^, all part of the 20 kbp core duplicon LCR16a ^44^. 1 additional match was found on chr18 and 11 matches on alternative chromosome scaffolds. The LCR16a region, also known as *morpheus* gene cluster, encodes variants of the nuclear pore complex interacting protein (NPIP) gene. It is one of the most extreme cases of positive selection observed in primates ^44,45^. Members of the NPIP gene family can be subdivided in two subfamilies, NPIPA and NPIPB, encoding for structurally different proteins and with the NPIPB subfamily being exclusively expanded in chimpanzee, human, and gorilla ^45,47^. In the majority of instances, PIPPI sORF is included in the last intron of these genes but this is not the case in a short splice variant of NPIPB5, in which the sORF is localizing to the 3’ UTR of the gene, and the long non-coding RNA PDXDC2P-NPIPB14P (Figure 2A). Interestingly, NPIPB5 seems to be the only paralogue of the NPIP gene family differentially expressed in the brain cortical tissue ^48^. Despite a high degree of similarity in the first 51 amino acids, the NPIPB14P locus encodes for a longer isoform of the microproteins, hereafter referred to as PIPPI-L. When we exogenously expressed the two GFP-tagged microproteins in A375 cells we noticed that PIPPI was more stable than PIPPI-L, therefore we decided to focus on this isoform for further characterization of the protein (Supplementary Figure 2A). To gain information on the endogenous expression and regulation of PIPPI, we raised a rabbit antiserum targeting a unique amino acid sequence present in this microprotein. Although we could affinity-purify antibody to detect exogenously expressed PIPPI-GFP (e.g. Supplementary Figure 2A), we were unable to detect endogenous PIPPI in parental A375, and the antibody showed various unspecific bands. This could be due to the low expression or absence of PIPPI in A375 cells, but also to the technical challenges, such as transferring the microprotein to a western blot membrane. In fact, using immunoblotting we failed to detect the exogenous expression of untagged PIPPI despite the microprotein being detectable by mass spectrometry (Figure 2E), suggesting that the untagged PIPPI was present but undetectable using our antibody.

**Figure 2:**
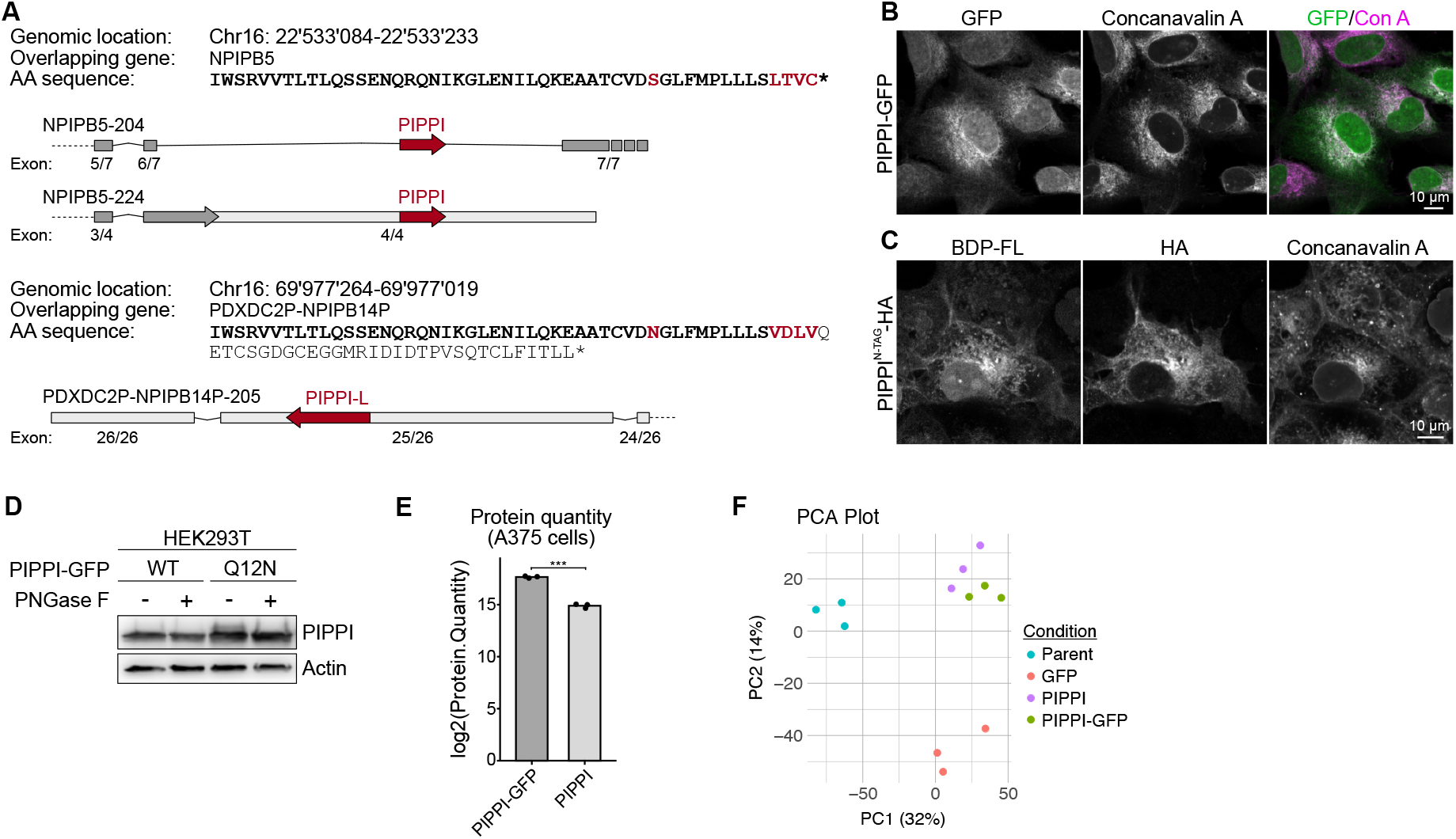
PIPPI is a novel microprotein localizing to the endoplasmic reticulum. A) Schema presenting the genomic coordinates and the organization of two selected loci encoding for PIPPI. B) Confocal images of RPE-1 cells stably expressing a PIPPI-GFP transgene. Concavalin A (Con A) staining was included to visualize the endoplasmic reticulum. C) STELLA tag was used to define the subcellular localization of PIPPI-HA in COS-7 cells. The BODIPY-FL-tetrazine dye (BDP-FL) was clicked to a ncAA amino acid incorporated in PIPPI-HA sequence. The endoplasmic reticulum is stained using Concavalin A. D) PIPPI-GFP constructs engineered to allow N-glycosylation were transiently expressed in HEK293T cells. After lysis, samples were treated with the recombinant amidase PNGase F and analysed by immunoblotting. E) Mass spectrometry was used to quantify the abundance of PIPPI and PIPPI-GFP in A375 cells. F) Plot presenting the principal component analysis (PCA) performed on the whole proteome data obtained from parental, GFP-, PIPPI and PIPPI-GFP- expressing A375 cells.

To characterize the cellular role of PIPPI, we overexpressed a tagged version of the microprotein in different cell lines. At this stage, we decided to work with different tags to ensure that these, due to their sizes and biophysical properties, would not affect the subcellular localization and function of the microprotein. A C-terminal HA-tag fusion, PIPPI- HA, appeared to be particularly lowly expressed and/or unstable, since we were not able to detect the tagged microprotein in transiently transfected (Supplementary Figure 2B) or stable cell lines (not shown) by immunoblotting. This limitation could be circumvented with the use of STELLA tag ^49^ that, possibly thanks to increased expression of the Ub-PIPPI-HA fusion protein, allowed the accumulation of a detectable pool of the microprotein both in HEK293T cells (Supplementary Figure 2B) and COS-7 cells (Figure 1C). Using STELLA tag, we could also show that in COS-7 cells PIPPI was enriched in the endoplasmic reticulum (ER), as highlighted by the good degree of colocalization with the lectin Concanavalin A, a well-established marker of this organelle (Figure 2C and Supplementary Figure 2C). This observation was confirmed in a panel of human cell lines stably overexpressing PIPPI-GFP (Figure 2B and Supplementary Figure 2D-E). To clarify whether PIPPI was associated to the cytoplasmic side of the ER membrane or was present within the ER lumen, we engineered a glycosylation site in PIPPI (PIPPI^Q12N^-GFP). When transiently expressed in HEK293T cells, this construct displayed an additional PIPPI band that migrated slower in SDS-PAGE and could be reverted by treatment with the de-glycosylase enzyme PNGase F (Figure 2D). Altogether, these results suggest that PIPPI is localizing within the glycosylation-supporting environment of the ER lumen ^50^, despite the fact that the microprotein does not possess a canonical ER localization sequence.

To verify that untagged PIPPI was present in A375 stable cell lines and assess how much overexpression of the microprotein impacts on cells, we performed a whole-proteome mass spectrometry (MS) analysis of parental, GFP-, PIPPI-, and PIPPI-GFP cells. While we were not able detect endogenous PIPPI-specific peptides in the parental cell line, MS showed that the microprotein was expressed in PIPPI and PIPPI-GFP stable cell lines and PIPPI levels were higher when the GFP tag was included (Figure 2E). Overall, overexpression of PIPPI or PIPPI-GFP only cased a small number of alteration to the proteome of A375 cells, and no particular pathway appeared to be up- or downregulated (Supplementary Figure 2F, Supplementary Table 3). Subjecting the MS results to a principal component analysis (PCA) highlighted that PIPPI and PIPPI-GFP were eliciting similar alterations. This suggested that appending a GFP tag to the microprotein sequence did not largely influence the cellular effects of PIPPI’s overexpression. Overall, we could show that when exogenously expressed in cells, PIPPI localizes in the lumen of the ER.

### The microprotein PIPPI enables cells to overcome stress

Based on the results of our initial screen, we hypothesized that PIPPI could be involved in the repair of DNA lesions and, in order to elucidate the role of PIPPI in the DNA damage response (DDR), we performed growth competition assays upon treatment with a panel of genotoxic agents. At this stage, we treated cells with either camptothecin, etoposide, cisplatin and oxygen peroxide to induce different types of DNA damage, but none of these treatments recapitulated the phenotype observed upon 6-TG administration. However, it is also known that 6-TG treatment can result in the activation of the unfolded protein response (UPR), a stress-response pathway that cells activate to relieve the accumulation of misfolded proteins in the ER ^51–53^. To assess if PIPPI modulated the UPR, parental and PIPPI-expressing A375 cells were treated for 5 hours with either 6-TG or tunicamycin (TM), a compound inducing the unfolded protein response (UPR) by inhibiting the first step of N-linked glycosylation ^54^. Stabilization of ATF4, a central player of the UPR, was used as a proxy for the activation of the pathway and confirmed that both treatments cause ER stress in A375 cells (Figure 3A). We further performed growth competition assays, in which cells were treated with a sub-lethal dose of TM. Confirming the involvement of PIPPI in an ER stress response pathway, PIPPI- expressing cells were able to grow better than their parental counterparts after either a short (5 hours) or long (24 hours) treatment with TM (Figure 2B). To better understand the effects of PIPPI on the cells’ ability to deal with misfolded proteins, we assessed the induction of some key components of the UPR. The chaperone BiP/HSPA5 plays a critical role in sensing the accumulation of misfolded proteins in the ER and, upon binding to unfolded proteins, it releases for activation the ER transmembrane transducers ATF6, IRE1 and PERK, which orchestrate downstream signaling events ^55^. ATF6 and IRE1 are transcription factors that promote the upregulation of ER chaperones, such as BiP itself, and folding enzymes, such as PDI. Differently, the PKR-like ER kinase PERK is one of the four kinases constituting the integrated stress response (ISR) and upon exposure to stress it can phosphorylate and inactivate the eukaryotic initiation factor 2 alpha (eIF2α) ^56^. As a result of eIF2α phosphorylation, the rate of translation initiation is decreased to reduce general protein synthesis and thereby alleviating the situation of stress. Under these conditions, a subset of mRNAs, which include the one encoding for activating transcription 4 (ATF4), are selectively translated to promote recovery and survival ^57^. Finally, if the stress situation is not promptly resolved, prolonged activation of the UPR results in the up-regulation of the C/EBP Homologous Protein (CHOP/DDIT3) transcription factor, which initiates apoptotic cell death^37^. Interestingly, despite both cell lines displayed a similar upregulation of BiP, indicating that they were experiencing the same level of ER stress, PIPPI-expressing cells had a dampened downstream signaling resulting in a weaker stabilization of both transcription factors ATF4 and CHOP (Figure 3C). Importantly, the same phenotype could also be recapitulated in A375 cells either expressing PIPPI-GFP or PIPPI-L (Supplementary Figure 3B-C). To confirm this observation using an independent approach, we engineered A375 to express a fluorescent reporter for ATF4 induction and imaged the living cells for 24 hours after treatment with TM ^58^. 2 hours after TM treatment, the levels of ATF4 started increasing in both parental and PIPPI- expressing cells but from the beginning the PIPPI curve was less steep than the parental one. Moreover, ATF4 levels plateaued earlier in PIPPI cells, corroborating the initial observation that these cells have a dampened activation of the UPR after exposure to ER stress. Finally, experiments performed in human U2OS osteosarcoma cells confirmed that the dampened activation of the UPR is not an A375-specific phenotype but can be recapitulated in cells of different origin (Supplementary Figure 3D-E). Summarizing, PIPPI-expressing cells have a dampened response when experiencing ER stress and, as a consequence, these cells have a better chance than parental cells to survive and proliferate under stress conditions.

**Figure 3:**
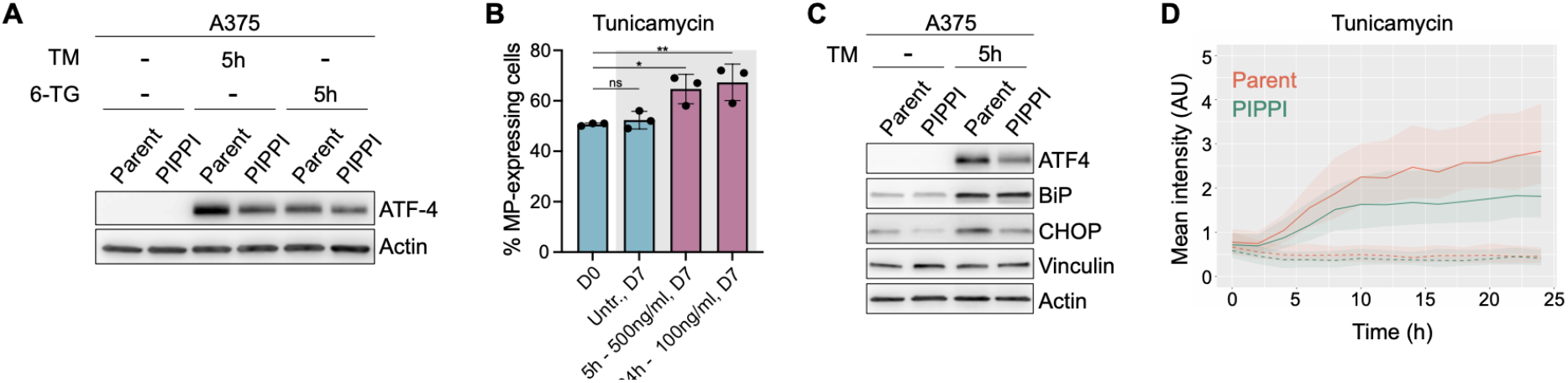
PIPPI is involved in the ER stress response. A) Parental and PIPPI-expressing A375 cells were treated for 5h with either 500 ng/ml Tunicamycin or 5µM 6-thioguanine. The level of ATF4 induction was assessed by immunoblotting. B) Bar chart presenting the results of the growth competition assays performed to assess the effects of tunicamycin on cellular proliferation of parental versus PIPPI-expressing cells. Height of the bars represents the fraction of PIPPI-expressing cells present in the total cell population. Values were averaged based on 3 independent biological replicates (black circles). p-values were calculated using unpaired Student’s t-test (ns = p > 0.05; * = p < 0.05, ** = p < 0.01). C) Parental and PIPPI-expressing A375 cells were treated with 500 ng/ml Tunicamycin for 5 hours. After lysis, the level of different proteins involved in the ER stress response were assessed by immunoblotting. D) Parental and PIPPI-expressing A375 cells, engineered to express a fluorescent ATF4 reporter, were treated with 500 ng/ml Tunicamycin and imaged every 30 minutes over the course of 24 hours. Wells with untreated cells were included and are reported in the graph as dashed lines. Plotted are the median, first quartile and third quartile values that were calculated using the mean intensity of at least 300 cells.

### Dissecting PIPPI’s interactome

To identify possible interaction partners of PIPPI, we performed co-immunoprecipitation (co-IP) experiments in which GFP-Trap was used to enrich proteins in parental, GFP- and PIPPI- GFP-expressing cells, before performing liquid-chromatography tandem mass-spectrometry (LC MS/MS). With this approach, we identified several ER-resident (BiP/HSPA5, P4HB, DNAJC10, ERLIN1) or ER-golgi trafficking proteins (ERp44, TMED10, TMED9) that were significantly enriched in the PIPPI-GFP sample in U2OS cells (Figure 4A-B, Supplementary Table 4). STRING network analysis of the top 10 putative PIPPI interactors highlighted proteins belonging to biological processes significantly enriched in the network. For example, blue nodes indicate proteins involved in the “response to ER stress” (FDR = 0.00054) and green nodes correspond to proteins related to “protein folding in the ER” (FDR = 0.00054). In addition to this, nodes associated with cellular components significantly enriched in the network, in this case the “ER-Golgi intermediate compartment”, are highlighted in red (FDR = 7.85e^-6^). Importantly, some of the identified proteins (i.e. ERp44, P4HB) were also found to be enriched in similar experiments performed in a different cellular background (A375 cells, Supplementary Figure 4A, Supplementary Table 4) and the interaction was not mediated by the presence of GFP (Supplementary Figure 4B). In follow up experiments, we validate the interaction of PIPPI-GFP with BiP and the Endoplasmic Reticulum Protein 44 (ERp44) (Figure 4C-D). BiP is a chaperone involved in the folding and assembly of proteins in the ER and its main function is to recognize and bind unfolded proteins in the ER. Hence, we were not able to distinguish if PIPPI-GFP is a folding client of BiP or binds BiP in another way. BiP protein levels increase when unfolded proteins accumulated in the ER, but we did not observe such effect by PIPPI expression, corroborating that PIPPI itself does not induce or increase protein folding stress in the ER (e.g. Figure 3C, Supplementary Figure 3C). ERp44 is a member of the protein disulfide isomerase (PDI) family of ER proteins, which principally function as a pH- and Zinc-dependent chaperone along the secretory pathway. In fact, ERp44 cycles between the ER and the Golgi, controlling the secretion of disulfide-linked oligomeric proteins and ensuring the correct localization of ER enzymes, lacking localization signals ^59^. Besides these canonical roles, ERp44 has been shown to interact and regulate a brain-specific subtype of the Inositol 1,4,5-trisphosphate receptors (IP3Rs), which are intracellular channels controlling calcium release from the ER ^60^. Finally, we also confirmed the interaction with the TMED10 and ERdj5/DNAJC10, but these two proteins are only weakly detectable in co-IP experiments (Figure 4E). TMED10 is a type I membrane protein which localizes to the plasma membrane and Golgi and is involved in protein trafficking ^61^. More recently, it has also been shown that TMED10 can mediate the uptake of leaderless secretory proteins into the ER-Golgi intermediate compartment (ERGIC), favoring their secretion via an unconventional protein secretion (UPS) pathway ^62^. ERdj5 is an ER disulfide reductase capable of both promoting correct folding of proteins by removing non-native disulfide bonds, but also for initiating the degradation of misfolded proteins ^63,64^.

**Figure 4:**
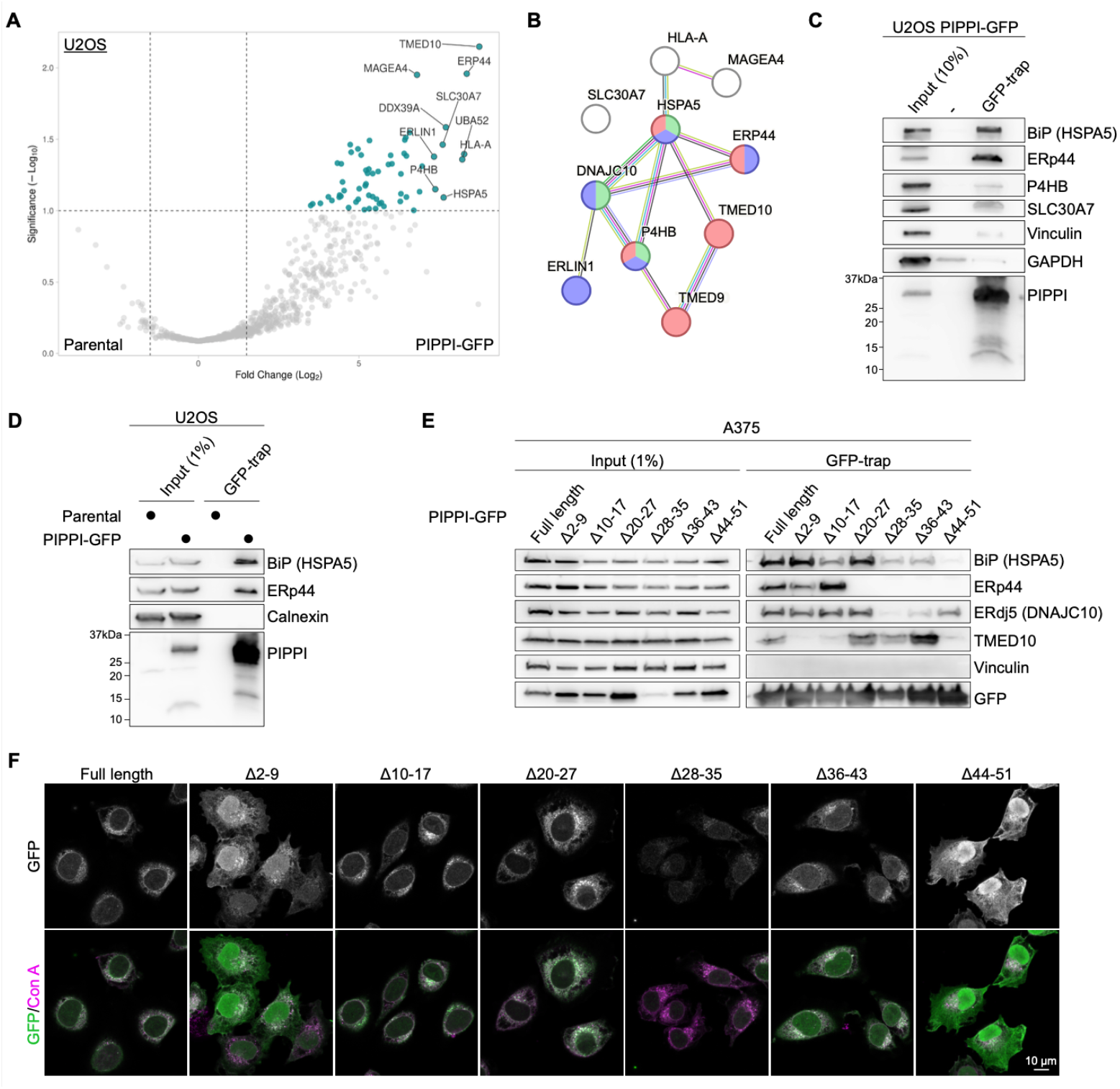
Identification of PIPPI interaction partners. A) Volcano plot of co-immunoprecipitation LC-MS/MS (co-IP/MS) experiments performed with parental and PIPPI-GFP-expressing U2OS cells. Experiment was conducted in two biological replicates, which were each analyzed three times by mass-spectrometry. Thresholds are set at Log2FC = 2 and DEP adjusted p-value = 0.1 (dashed lines). Proteins enriched in PIPPI- GFP lysates are clustering in the right half of the plot and the top ten proteins (assessed by Manhattan distance) are labelled with their names. B) STRING network of the putative top 10 PIPPI interactors identified by co-IP/MS. Nodes belonging to biological processes significantly enriched in the network are highlighted in blue (Response to ER stress, FDR = 0.00054) and in green (Protein folding in the ER, FDR = 0.00054). Nodes associated to cellular components significantly enriched in the network are highlighted in red (ER-Golgi intermediate compartment, FDR = 7.85e^-6^). C) U2OS-PIPPI-GFP lysates were immunoprecipitated using a GFP-trap and analysed by immunoblotting to validate the co-IP/MS results. D) Lysates obtained from parental and PIPPI-GFP-expressing U2OS cells were immunoprecipitated using a GFP-trap and analysed by immunoblotting. E) Lysates from A375 cells stably expressing different deletion mutants of PIPPI were immunoprecipitated using a GFP-trap and analysed by immunoblotting to identify regions in PIPPI responsible for protein-protein interactions. F) Confocal images of A375 cells stably expressing a panel of PIPPI-GFP truncations. Concavalin A (Con A) staining was included to visualize the endoplasmic reticulum.

To better understand the relationship between PIPPI and its interaction partners, we generated a panel of cell lines expressing PIPPI-GFP truncations (Supplementary Figure 4C) and analysed them by co-IP and confocal microscopy. The most striking phenotype was observed for PIPPI^Δ44–51^, lacking its C-terminus, resulting in the loss of almost all interaction partners (Figure 4E) and a pan-cellular localization (Figure 4F). A diffuse localization was also observed for the N-terminal mutant PIPPI^Δ2–9^, despite this mutant seemed to retain some of the interaction partners (Figure 4E). Interestingly, both of these PIPPI mutants lost their interaction with TMED10. Thus, we hypothesize that in normal conditions, the Golgi complex- localizing protein channel TMED10 is shuttling PIPPI into the lumen of the secretory pathway, where it is retained principally thanks to the action of ERp44. Another interesting phenotype could be observed for PIPPI^Δ28–35^, which appeared to be particularly unstable (Figure 4E-F). Finally, the Δ10-17, Δ20-27 and Δ36-43 mutants appeared to retain their capability of localizing in the ER and Golgi complex, but their distribution within the compartments could be affected by their differential ability to interact with ERp44, BiP and ERdj5. Interestingly, mutants Δ20-27 and Δ36-43 seemed to be able to reach the ER, as judged by their capability to interact with BiP and ERdj5, despite not interacting with ERp44 (Figure 4E). This indicates that there is another factor able to relocate PIPPI from the Golgi to the ER. Further, mutants of the C-terminal domain of PIPPI, where the only two cysteines of PIPPI are located, interact less promptly with ERdj5. To summarize, we showed that PIPPI is interacting with different proteins within the secretory compartment, predominantly with ERp44 and BiP, and tentatively delineate the role of each interactor for PIPPI localization and function.

### The ability to interact with ERp44 is a prerequisite for PIPPI’s function

To further validate the interaction between PIPPI and ERp44, we performed reverse co-IP by immunoprecipitating HA-ERp44 in lysates obtained from PIPPI-GFP A375 cells engineered to co-express HA-ERp44 (Supplementary Figure 5A). To corroborate this result, we performed the same kind of immunoprecipitation experiment after co-overexpressing ERp44 and PIPPI- GFP in HEK293T cells (Figure 5A). In this experiment, we included the ERp44^C29V^ mutant which is unable to interact with his client proteins ^65^. The mutated cysteine C29, which is part of the N-terminal thioredoxin-like domain of ERp44 (CRFS), is in fact necessary to form mixed disulfide bonds with client proteins ^65^. Interestingly, the C29V mutant of ERp44 was not abrogating the interaction with PIPPI, suggesting that PIPPI is not a canonical client protein of ERp44 (Figure 5A). To clarify the importance of (mixed) disulfide bonds for PIPPI function and encouraged by the observation that PIPPI^Δ28–35^ and PIPPI^Δ44–51^ severely affect the microprotein’s stability and localization, we decided to point-mutate to alanine the two cysteines present in PIPPI (C35A and/or C51A). The double mutant PIPPI^C35A-C51A^ appeared to be particularly unstable, but the single point mutants could be expressed in cells and were slightly more stable than the *wild type* microprotein, suggesting that the two cysteine forming an intramolecular disulfide bonds is not a prerequisite for proper folding and stabilization of the protein. When we tested how the cysteine mutants affected the interactome of PIPPI, we noticed that both point mutants completely abrogated the interaction with ERp44 and increased the fraction of the protein that remains bound to TMED10 (Figure 5B). Similar to what observed with the truncation mutants, inhibiting the ability of PIPPI to interact with ERp44 decreased the amount of the microprotein that reaches the ER, as suggested by the lower amount of PIPPI binding to BiP (Figure 5B). Finally, we investigated how these two PIPPI mutants influenced cell proliferation upon treatment with tunicamycin and we observed that overexpression of neither of them was helping cells coping with ER stress to the extent measured with the *wild type* microprotein (Figure 5C). In line with this, the dampened activation of the UPR observed in PIPPI cells was less accentuated in cells expressing the one or the other cysteine mutant, suggesting that interaction with ERp44 is critical for PIPPI functionality (Supplementary Figure 5B).

**Figure 5:**
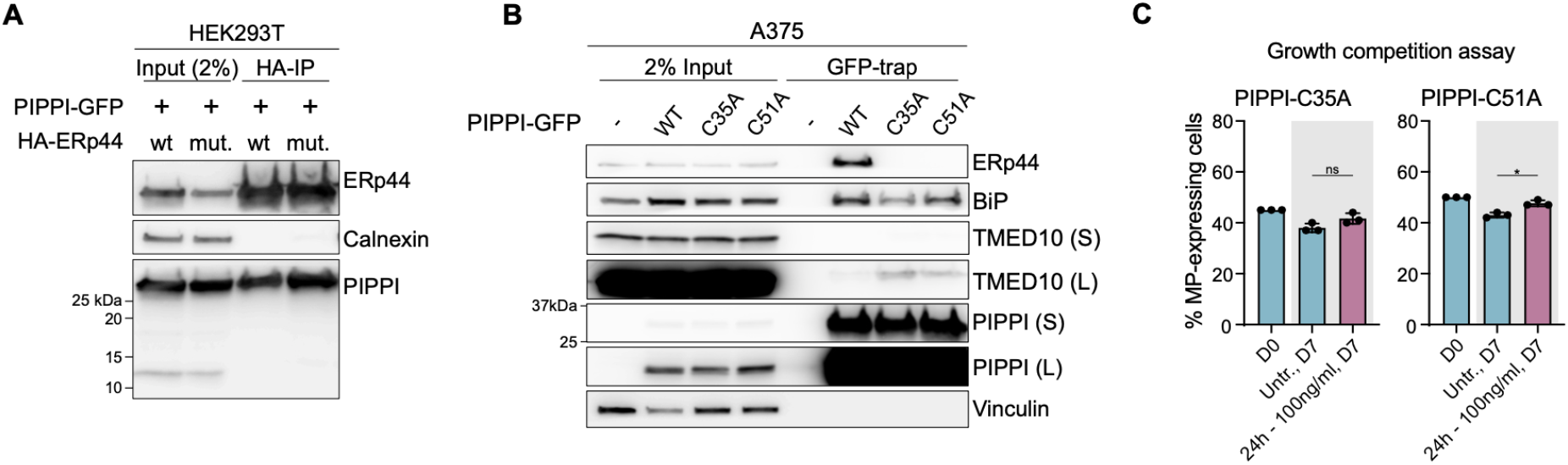
PIPPI exerts his protective function through the protein disulfide isomerase ERp44. A) PIPPI-GFP was transiently co-overexpressed with either wild-type HA-ERp44 or a *CRSF mutant* of HA-ERp44 in HEK293T cells. Lysates were subjected to immunoprecipitation using HA-beads and analysed by immunoblotting. B) Lysates obtained from parental, *wild type* PIPPI-GFP-, *C35A* PIPPI-GFP-, and *C51A* PIPPI-GFP-expressing A375 cells were subjected to immunoprecipitation using a GFP-trap and analysed by immunoblotting. (S) indicates a short exposure, whereas (L) indicates a long exposure of the same antibody. C) Bar chart presenting the results of the growth competition assays performed to assess the effects of tunicamycin on the proliferation of *C35A* PIPPI-GFP- and *C51A* PIPPI-GFP-expressing A375 cells. Height of the bars represents the fraction of PIPPI-expressing cells present in the total cell population. Values were averaged based on 3 independent biological replicates (black circles). At least 10’000 cells were measured for each sample.

## Discussion

In this manuscript we presented a novel approach for screening a large number (>10’000) of non-canonical sORFs, based on the pooled overexpression of synthetic sORFs. An arrayed sORFs screen to assess translation and stability of microproteins have already been performed recently ^66^, but the presented approach is suitable for the investigation of a larger number of coding sequences without the need of specialized instrumentation (i.e. liquid handling robots). In a proof-of-concept screen performed to uncover novel factors promoting cell survival upon exposure to 6-thioguanine, we identified the known microprotein altDDIT3 and the novel, uncharacterized microprotein PIPPI. PIPPI is encoded within a highly duplicated region of the human genome, which also entails the understudied *nuclear pore complex interacting protein* (NPIP) gene family ^44,67^. Based on our data, the microprotein PIPPI seems to be functionally unrelated from the NPIP gene products ^47^. Interestingly, these genomic regions are undergoing rapid positive selection in primates ^44^, suggesting that these evolutionary young proteins could perform specialized functions in primate-specific tissues and organs. A limitation of our study is that we were not able to define conditions under which PIPPI is expressed, and we could not delineate which (one or more) instances of the sORF could produce the microprotein. While PIPPI was first suggested on the basis of proteomics evidence ^43^, we were not able to detect and study endogenous PIPPI with an antibody we raised. Hence, we worked with the exogenously expressed microprotein to define its localization and interactome to set the basis for further characterization of PIPPI. Interestingly, PIPPI appeared to be localizing in the ER lumen, despite being devoid of canonical signal peptides or ER localization motifs. We hypothesized that this occurs thanks to the action of the trafficking protein TMED10, which can orchestrate the secretion of a spectrum of cargos upon facilitating their uptake in the ER-Golgi intermediate compartment (ERGIC) ^62^. From here, PIPPI can be shuttled to the ER via its interaction with the PDI-homologue chaperone ERp44 ^59^. Once in the ER, we observed that PIPPI has the ability to positively affect the cellular response to ER stress, allowing cells to proliferate better, and partially escape apoptosis, under these conditions. Challenges remain to understand how and when specific sORFs are used by an organism, especially in the case of recently evolved sORFs, whose expression and function could be extremely narrow by cell type or condition. As an attenuator of the ISR, PIPPI has a general pro-survival function that could be invoked by cells under specific stress conditions, which we yet have to fully understand. Nevertheless our study highlights the functional potential of the vast and largely uncharacterized human sORF translatome.

## AUTHOR CONTRIBUTIONS

Conceptualization: LL, SJE. Methodology: LL, CN, DS, MH, IP, SJE. Investigation: LL, AS, MH, LB, DS, CN, RS, IP, SJE. Formal analysis: LL, DH, IP,, SJE. Writing-original draft: LL, SJE. Writing-review and editing: LL, DS, CN, AS MH, IP, SJE. Funding acquisition: SJE. Resources: SJE. Supervision: SJE.

## ACKNOWLEDGEMENTS

S.J.E. acknowledges funding by the Karolinska Institutet SFO for Molecular Biosciences, Vetenskapsrådet (2015-04815, 2020-04313), H2020 ERC Starting Grant (715024 RAPID), Åke Wibergs Stiftelse (M15-0275), Cancerfonden (22 2354 Pj), the Ming Wai Lau Center for Reparative Medicine, the Ragnar Söderbergs Stiftelse, Stiftelsen för Strategisk Forskning (FFL7); and the Knut och Alice Wallenbergs Stiftelse (2010–0215). DS was supported by a Boehringer Ingelheim Fonds PhD fellowship. Sequencing analyses were processed on the Uppsala Multidisciplinary Center for Advanced Computational Science (UPPMAX) provided by the National Academic Infrastructure for Supercomputing in Sweden (NAISS) under projects NAISS 2023/6-19, NAISS 2023/22-84, SNIC 2022/6-14. NAISS is funded by the Swedish Research Council through grant agreement no. 2022-06725. We thank Prof. Iris Finkemeier and Paulina Heinkow for mass spectrometry service. We thank the BIC facility for enabling usage of their AiryScan microscopy, the Fernandez Capetillo group for allowing us to use their In Cell microscope, the Bartek and Lemmens groups for access to the Nikon Eclipse Ti2.

## Materials and methods

### Cell culture

A375, U2OS, HEK293T, and COS-7 cell lines were maintained in Dulbecco’s Modified Eagle Medium (DMEM), containing high glucose, GlutaMAX^TM^ and pyruvate (Gibco). hTERT RPE-1 cells were maintained in Dulbecco’s Modified Eagle Medium/Nutrient Mixture F-12 (DMEM/F- 12), containing high glucose, GlutaMAX^TM^ and pyruvate (Gibco). For all cell lines, medium was supplemented with 10% fetal bovine serum (FBS, Sigma-Aldrich). All cell lines were cultured in an ambient-controlled incubator at 37°C, 5% O2 and 5% CO2. A375 and COS-7 cells were purchased from Sigma-Aldrich. HEK293T cells were a gift from Prof. T. Helleday, whereas U2OS, and RPE-1 cells were a gift from Prof. J. Bartek.

### DNA constructs

sORF sequences were synthesized by Twist Biosciences and cloned in the destination vectors by In-Fusion Cloning (Takara Bio) or Golden Gate cloning. All DNA constructs were verified by Sanger sequencing.

| ID | Name/description | Backbone | Origin |
| --- | --- | --- | --- |
| CD532A-2 | pCDH_EF1a_MCS_IRES_Puro | pCDH | System Biosciences |
| L300 | pCDH_EF1a_MCS_IRES_Puro_SapI_removed | CD532-A | This study |
| L301 | pCDH_EF1a_2xSapI_A_IRES_Puro | L300 | This study |
| L302 | pCDH_EF1a_2x SapI_C_IRES_Puro | L300 | This study |
| L303 | pCDH_EF1a_2x SapI_G_IRES_Puro | L300 | This study |
| L309 | pCDH_EF1a_All-sORFs_IRES_Puro | L301/L302/L303 | This study |
| L320 | pCDH_EF1a_altDDIT3_IRES_Puro | L302 | This study |
| #12260 | psPAX2 (2<sup>nd</sup> generation lentiviral packaging plasmid) |  | Addgene |
| #12259 | pMD2.G (VSV-G envelope expressing plasmid) |  | Addgene |
| L321 | pCDH_EF1a_altDDIT3-GFP_IRES_Puro | L327 | This study |
| L325 | pAS1_EF1a_PIPPI-HA_IRES_Puro | E404 (#140009) | This study |
| K106 | pAS1_4x7SKPyIT_EF1_Ub-*PIPPI | K038 (#162801) | This study |
| K110 | pAS1_4x7SKPyIT_EF1_Ub-*PIPPI-HA | K039 (#162802) | This study |
| K111 | pAS1_4x7SKPyIT_EF1_Ub-PIPPI-HA | K039 (#162802) | This study |
| L327 | pCDH_EF1a_2x SapI_C-GFP_IRES_Puro | L302 | This study |
| L329 | pCDH_EF1a_PIPPI_IRES_Puro | L302 | This study |
| L356 | pCDH_EF1a_GFP_IRES_Puro | L313 | This study |
| L330 | pCDH_EF1a_PIPPI-GFP_IRES_Puro | L327 | This study |
| L344 | pCDH_EF1a_PIPPI-L-GFP_IRES_Puro | L327 | This study |
| L346 | pCDH_EF1a_2x SapI_C-GFP_IRES_BSD | L327 | This study |
| L348 | pCDH_EF1a_PIPPI-GFP_IRES_BSD | L346 | This study |
| #115969 | pLVX-ATF4 mScarlet NLS | pLVX-Puro | Addgene |
| L363 | pCDH_EF1a_PIPPI-L_IRES_Puro | L302 | This study |
| L381 | pCDH_EF1a_PIPPI-del2-9-GFP_IRES_Puro | L327 | This study |
| L382 | pCDH_EF1a_PIPPI-del10-17-GFP_IRES_Puro | L327 | This study |
| L383 | pCDH_EF1a_PIPPI-del20-27-GFP_IRES_Puro | L327 | This study |
| L384 | pCDH_EF1a_PIPPI-del28-35-GFP_IRES_Puro | L327 | This study |
| L385 | pCDH_EF1a_PIPPI-del36-43-GFP_IRES_Puro | L327 | This study |
| L386 | pCDH_EF1a_PIPPI-del44-51-GFP_IRES_Puro | L327 | This study |
| A014 | pCDH_EF1a_2x SapI_G_IRES_BSD | L303 | This study |
| L388 | pCDH_EF1a_HA-ERp44_IRES_BSD | A014 | This study |
| L389 | pCDH_EF1a_HA-ERp44_CRFS-mut_IRES_BSD | A014 | This study |
| L405 | pCDH_EF1a_PIPPI-C35A_IRES_Puro | L302 | This study |
| L406 | pCDH_EF1a_PIPPI-C35A-GFP_IRES_Puro | L327 | This study |
| L407 | pCDH_EF1a_PIPPI-C51A_IRES_Puro | L302 | This study |
| L408 | pCDH_EF1a_PIPPI-C51A-GFP_IRES_Puro | L327 | This study |
| L410 | pCDH_EF1a_PIPPI-Q12N-GFP_IRES_Puro | L327 | This study |

### Generation of stable expression cell lines using lentivirus

One day after seeding, HEK293T cells were forward co-transfected with a lentiviral transfer plasmid, the envelope plasmid pMD2.G (Addgene #12260) and the 2^nd^ generation packaging plasmid psPAX2 (Addgene #12259) using transIT^®^-LT1 (Mirus Bio) according to manufacturer’s instructions. After 24 hours, medium was replenished with fresh DMEM. Virus supernatant was collected 48 and 72 hours and, after filtration using a 0.45mm mixed cellulose esters syringe filter (Millipore), directly used to transduce target cells. The following concentrations of polybrene (Sigma-Aldrich), puromycin (Cayman Chemical), or blasticidin (InvivoGen) were used to either help infecting or select target cells: A375 (2µg/ml, 3µg/ml, 20µg/ml), U2OS (4µg/ml, 6µg/ml, -), hTERT RPE-1 (4µg/ml, -, 50µg/ml).

### sORF library design and production

Oligonucleotide pools were synthetized at TWIST bioscience. SapI sites surrounded the sORF sequences to enable cloning into the custom-modified pCDH_EF1_IRES_Puro lentiviral expression plasmid (System Biosciences, CD532A-2). Following golden gate cloning using SapI (NEB) and T4 ligase (NEB), the vector was electroporated into MegaX DH10BTM T1^R^ cells (Invitrogen). Bacteria were plated and selected overnight with 200mg/ml Carbenicillin. At this stage, a number of plates ensuring the maintenance of library diversity was used. Colonies were collected by scraping and pooled before extracting plasmid DNA using the QIAGEN Plasmid Plus Maxi Kit (Qiagen).

### Virus production for pooled screens

A concentrated lentivirus pool was produced at the VirusTech Core Facility of Karolinska Institutet. In short, HEK293T cells were co-transfected with the lentiviral expression plasmid pCDH_EF1-sORFs_IRES_Puro, the envelope plasmid pMD2.G (Addgene #12260) and the 2^nd^ generation packaging plasmid psPAX2 (Addgene #12259). pMD2.G and psPAX2 plasmids were gifts from Didier Trono. The media was exchanged after 16 hours and virus-containing supernatant was collected at 48- and 64-hours post transfection. After concentrating by double step centrifugation, the final titer was assessed by extracting and quantifying proviral DNA from transduced cells. The number of integrations into the host genome was calculated by normalizing the total number of provirus copies to a housekeeping gene (hALB).

### sORF screens

During screens, media was supplemented with 1% penicillin-streptomycin. A375 cells were transduced at day 0 with the lentiviral sORF-encoding library in two biological replicates at a low MOI (∼0.5). For each replicate, transduction was performed in presence of 2µg/ml polybrene (Sigma-Aldrich) and in enough cells to achieve a representation of at least 1000 cells per sORF. Two days after transduction, cells were re-seeded and selected with 1.5 mg/ml puromycin. At day 5, a fraction of cells was pelleted by centrifugation and frozen for genomic DNA extraction (timepoint 0, “T0”). The remaining cells were kept in puromycin and split in different arms for further “no drug” or drug treatments. After one day, puromycin selection was removed and treatment with 10 mM 6-thioguanine (Sigma-Aldrich) was started. Cells of the “no drug” control arm were maintained under puromycin selection for three days longer. Untreated cells were sub-cultured every 3-4 days until the final timepoint at day 21. Drug treatment was re-new every third day and cells were subcultured when and if needed. Screen was continued until cells repopulated the culture flasks (day 28). Throughout the screen, cells were collected or re-seeded in a number that would maintain a representation of at least 1000 cells per sORF.

### Genomic DNA extraction and sequencing

Genomic DNA (gDNA) was isolated using the QIAamp DNA Blood Midi Kit (Qiagen) accordingly to the manufacturer’s instructions. For PCR amplification of sORF sequences, gDNA was divided into 100 μL reactions such that each tube had at most 4 μg of gDNA. PCR reactions were performed using NEBNext^®^ Ultra^TM^ II Q5^®^ Master Mix (NEB). Afterwards, tubes were pooled per sample and purified using the QIAquick PCR Purification Kit (Qiagen). Custom-made primers containing the i5/i7 adapter sequences were synthesized at Integrated DNA Technologies (IDT) and incorporated during a second PCR step. PCR products were purified using QIAquick PCR Purification Kit (Qiagen) and sequenced on the Illumina NextSeq500 platform. Finally, the resulting Fastq files were aligned to our custom sORF library using kallisto v0.46.1 ^23^.

### Growth competition assays and flow cytometry

One day before treatment cell lines were seeded in a 1:1 ratio. This time point is considered “Day 0” of each assay and an aliquot of the cell mix was collected for flow cytometry analysis. On day 1, cells were treated with the indicated dose of 6-thioguanine (6-TG) or tunicamycin (Sigma-Aldrich) for either 5 or 24 hours (see figures for details). An untreated control well was included for each replica and, except the assay shown in Figure 4D, biological replicas were never performed in parallel. Throughout the duration of the assay, cells were visually assessed for viability and confluency, and split accordingly. On sampling days, cells were collected by trypsinization and a fraction of the suspension was fixed with 4% Formaldehyde for 10 min at room temperature. After washing, cells were resuspended in PBS and stored in the dark at 4°C until analyzing them with a Navios flow cytometer (Beckman Coulter). At least 10’000 living cells were measured for each sample. Long-term growth competition assays shown in supplementary figure 2 were performed as above but 6-TG was renewed every third day and cells were collected for flow cytometry every 7^th^ day. In this case, biological replicates were performed in parallel.

### Antibodies

| Target | Antibody name | Company | Application |
| --- | --- | --- | --- |
| ATF-4 | Anti-ATF-4 (D4B8) rabbit mAb, #11815 | Cell Signaling | WB (1:1000) |
| b-Actin | b-Actin (13E5) rabbit mAb, #4970 | Cell Signaling | WB (1:5000) |
| BiP | BiP (C50B12) rabbit mAb, #3177 | Cell Signaling | WB (1:1000) |
| Calnexin | Calnexin (C5C9) rabbit mAb, #2679 | Cell Signaling | WB (1:1000), IF (1:50) |
| CHOP | CHOP (L63F7) mouse mAb, #2895 | Cell Signaling | WB (1:500) |
| ERp44 | ERp44 rabbit Ab, #2886 | Cell Signaling | WB (1:500) |
| ERp44 | TXNDC4 (E-6) mouse mAb, sc-393687 | Santa Cruz | IP |
| GAPDH | GAPDH chicken Ab, AB2302 | Millipore | WB (1:10'000) |
| GFP | GFP (B-2) mouse mAb, sc-9996 | Santa Cruz | WB (1:5000) |
| GFP | GFP rabbit Ab, pabg-1 | ChromoTek | IF (1:800) |
| HA | HA-HRP high affinity Ab, 12013819001 | Roche | WB (1:5000) |
| HA | HA-Tag (F-7) mouse mAb, sc-7392w | Santa Cruz | IF (1:400) |
| P4HB | PDI (C81H6) rabbit mAb, #3501 | Cell Signaling | WB (1:1000) |
| SLC30A7 | SLC30A7 rabbit Ab, HPA018034 | Atlas Antibodies | WB (1:200) |
| Vinculin | Vinculin, rabbit Ab, #4650 | Cell Signaling | WB (1:1000) |

### Generation of PIPPI antibody

The PIPPI antibody was generated by *immunoGlobe GmbH.* In short, a rabbit was immunized using the 11-AA-long synthetic peptide CSENQRQNIKG, corresponding to PIPPI amino acids 14-23. Serum from the final bleed was affinity-purified using the High-Affinity Antibody Purification Kit (GenScript) and used for immunoblotting experiments (1:200-1:500 dilutions).

### Protein gels and western blotting

Cell lysates were prepared in RIPA buffer (50 mM Tris–HCl, pH 7.5, 1% NP-40, 0.25% sodium deoxycholate, 150 mM NaCl, 1 mM EDTA, and 0.1% SDS) supplemented with 1x cOmplete^TM^ Protease inhibitor cocktail (Roche). After sonication, the insoluble fraction was removed by centrifugation. Proteins were resolved by SDS–PAGE under denaturing conditions using 4- 20% Mini-PROTEAN^®^ TGX^TM^ precasted polyacrylamide gels (Biorad). Proteins were transferred to nitrocellulose membranes (Biorad). Immunoblots were performed using the appropriate primary antibodies and the relative HPR-coupled secondary antibodies (Biorad). Proteins were visualized on an ImageQuant^TM^ LAS500 Imager (GE Healthcare life Sciences) using the Immobilon Forte Western HRP substrate (Merck Millipore).

To visualize ncAA-containing proteins, lysates were incubated before denaturation in presence of 500nM SiR- tetrazine (Spirochrome) for 15 minutes at room temperature. After electrophorasis, the gel was imaged using an Amersham Imager 600 (GE Healthcare life Sciences).

### Amber suppression

HEK293T were forward co-transfected with a plasmid encoding the protein of interest and a plasmid encoding the *AF*variant of the *Methanosarcina mazei* pyrrolysine tRNA synthetase (pylRS-AF) using transIT^®^-LT1 (Mirus Bio) according to manufacturer’s instructions. COS-7 cells were forward transfected using Lipofectamine LTX™ with PLUS™ reagent (Invitrogen) according to manufacturer’s protocol. The non-canonical amino acid (ncAA) was added at the time of transfection and cells were either fixed after 24 hours for microscopy (COS-7) or lysed after 48 hours and analysed by immunoblotting (HEK293T). When cells were to be analysed by microscopy, the ncAA was withdrawn one hour before cell fixation. For all experiments axial *trans*-cyclooct-2-ene-l-lysine (TCO*K) was used as ncAA. TCO*K (SiChem, SC-8008) stock solution was prepared at 100 mM in 0.2 M NaOH/H2O, 15% DMSO and diluted to 50 µM right before use in the appropriate growth medium. At the end of the experiment, TCO*K-containing polypeptides were labeled using either Tetrazine-Silicon Rhodamine (tet-SiR, Spirochrome) or 6-Methyl-Tetrazine-BODIPY^®^-FL (me-tet-BDP-FL, Jena Bioscience). Both stocks were prepared in DMF and further diluted in either RIPA buffer (lysate labeling) or TBS-T (fixed cells labeling) for final use.

### Analysis of glycosylated proteins

Constructs were forward transfected in HEK293T cells using transIT^®^-LT1 (Mirus Bio) according to manufacturer’s instructions. After 24 hours, cells were lysed in RIPA buffer and, after sonication, the insoluble fraction was removed by centrifugation. Samples were then treated with PNGase F (NEB) according to manufacturer’s instructions. In short, lysates were denatured for 10 minutes in Denaturing Buffer at 100 °C. After cooling, samples were incubated for 60 minutes at 37°C in presence of 1% NP-40, GlycoBuffer 2 and PNGase F. Negative control samples were included. All samples were then analysed by SDS-PAGE gels and immunoblotting.

### Immunoprecipitation

Cell lysates were prepared from ca. 80% confluent T-175 flasks in RIPA buffer not containing SDS (50 mM Tris–HCl, pH 7.5, 1% NP-40, 0.25% sodium deoxycholate, 150 mM NaCl, 1 mM EDTA) supplemented with 1x cOmplete^TM^ Protease inhibitor cocktail (Roche). After allowing lysis for 15min rotating at 4°C, the insoluble fraction was removed by centrifugation. Supernatants were incubated for 4 hours rotating at 4°C with either 25ul of GFP-trap magnetic beads (ChromTek, pre-equilibrated in RIPA w/o SDS) or anti-HA magnetic beads (Pierce^TM^, pre-equilibrated in 0.05% TBS-T). In the case of the IP-LC MS/MS experiment in U2OS cells, RIPA was supplemented with 0.1% SDS for lysis. Before incubating the lysates with the magnetic beads, samples were diluted 1:10 in RIPA w/o SDS to lower the concentration of the detergent. After incubation with the lysates, beads were bound to a magnetic stand and, if not specified otherwise, washed three times with RIPA w/o SDS and once with ddH2O. Samples were shortly vortexed in-between washes. Finally, proteins were eluted from beads either by boiling in 2x Laemmli buffer (4% SDS, 20% glycerol, 120mM pH 6.8) for western blotting analysis or using 1% formic acid for LC-MS/MS analysis. Elution with 1% formic acid was performed twice and supernatants were pooled for downstream processing.

### Sample preparation for IP-mass spectrometry

After neutralization, samples were reduced and alkylated for 30min at room temperature with 5mM TCEP (Pierce^TM^) and 20mM chloroacetamide (Sigma) dissolved in 250mM Tris buffer pH 8.5. Afterwards, proteins were purified using SP3 beds. In short, Sera-Mag beads (Sigma-Aldrich) were pre-mixed 1:1 and added to the samples in a 1:25 ratio. An equal volume of absolute ethanol was added to the mixture to stimulate coupling of the proteins to beads. After 5min incubation at room temperature, beads were washed on a magnetic stand 3 times with 80% ethanol and then resuspended in 50mM TEAB (Sigma-Aldrich). 1µg trypsin (Pierce^TM^) was added to each sample and digestion was allowed to occur o.N. at 37°C under gentle agitation. After immobilizing beads on a magnetic stand, supernatants were collected. Bound peptides were then eluted using 2% Acetonitrile (Sigma-Aldrich) in 50mM TEAB (Sigma-Aldrich) and pooled with the collected supernatants. Samples were completely dried using a SpeedVac and finally resuspended in solvent B (80% Acetonitrile, 0.1% formaldehyde) for LC-MS/MS analysis.

### IP-Mass spectrometry (LC-MS/MS)

LC-MS/MS was provided on a service basis by the Finkemeier lab (Münster University). Samples were analyzed using an EASY-nLC 1200 (Thermo Fisher Scientific) coupled to an Exploris 480 mass spectrometer (Thermo Fisher Scientific). Separation of peptides was performed on 20 cm frit-less silica emitters (CoAnn Technologies, 0.75 μm inner diameter), packed in-house with reversed-phase ReproSil-Pur C18 AQ 1.9 μm resin (Dr. Maisch). The column was constantly kept at 50 °C. Peptides were eluted in 115 min applying a segmented linear gradient of 0 % to 98 % solvent B (solvent A 0 % ACN, 0.1 % FA; solvent B 80 % ACN, 0.1 % FA) at a flow-rate of 300 nL/min. Mass spectra were acquired in data- dependent acquisition mode. MS1 scans were acquired at an Orbitrap Resolution of 120,000 with a Scan Range (m/z) of 380-1500, a maximum injection time of 100 ms and a Normalised AGC Target of 300 %. For fragmentation only precursors with charge states 2-6 were considered. Up to 20 Dependent Scans were taken. For dynamic exclusion, the exclusion duration was set to 40 sec and a mass tolerance of +/- 10 ppm. The Isolation Window was set to 1.6 m/z with no offset. A normalised collision energy of 30 was used. MS2 scans were taken at an Orbitrap Resolution of 15,000, with a fixed First Mass (m/z) = 120. Maximum injection time was 22 ms and the normalised AGC Target 50 %.

### Immunofluorescence microscopy

Cells grown in 18 well μ-Slides (ibidi) were fixed with 4% formaldehyde for 10 min at room temperature and permeabilized for 15 min with 0.1% (v/v) triton/PBS. After washing with TBS supplemented with 0.1% Tween-20 (TBS-T), cells were blocked with 2% BSA in TBS-T for 1 hour and then incubated overnight at 4°C with primary antibodies.In amber suppression experiments, cells were click-labeled for 60 min with 500 nM 6-Methyl-tetrazine-BODIPY-FL in blocking buffer (2% BSA in TBS-T) and washed three times with TBS-T prior to the incubation with primary antibodies. After washing with TBS-T, cells were stained with Alexa Fluor-conjugated secondary antibodies (Life Technologies) for 60 min at room temperature and counterstained with 1 mg/ml DAPI (Sigma-Aldrich) or with 10 μg/ml AF647-conjugated Concanavalin A (Invitrogen). After washing, cells were imaged on a Zeiss LSM880 confocal laser scanning microscope using a 63x/1.4 oil immersion objective. Images were processed and prepared for publication using Fiji ^24^.

### Live cell microscopy

One day before the experiment, A375 cells expressing the ATF4-mScarlet-NLS reporter were seeded in 18 well μ-Slides (ibidi) and cultured overnight in DMEM supplemented as detailed above. One hour before starting the experiment, medium was exchanged to Leibovitz’s L-15 Medium (no phenol red, Gibco) supplemented with 10% FBS (Sigma-Aldrich), 4.5 g/L glucose (Sigma-Aldrich) and 1mM Sodium Pyruvate (Gibco). Right before imaging, Tunicamycin (500ng/ml) were added to some of the wells. Cells were imaged over 24 hours at 2-hour intervals using a Nikon eclipse Ti2 inverted widefield microscope equipped with a heated imaging chamber. Images were acquired with a 20×/0.75 air objective and, post-acquisition, quantified using CellProfiler ^25^. Plots included in the publications were prepared using ggplot2 v3.3.5 ^26^ run on RStudio v2022.02.03 ^27^.

### MS data analysis

The raw data was analysed using MaxQuant Version 1.6.3.4 ^28^ and searched against the Uniprot protein database (Human all 2017/11) ^29^, the PIPPI-GFP sequence and the CRAPome database ^30^. MaxQuant default settings were used with the two options *Match between runs* and *LFQ intensity reporting* activated. The resulting LFQ intensities were then analysed using the Differential Enrichment analysis of Proteomics data (DEP) package 1.14.0 ^31^ on RStudio 2022.02.03. After removing common contaminants, imputation was performed using the “MinProb” method without a prior “DEP normalization” of the data. Proteins were then ranked according to their LFC enrichment over parental control and DEP p-value. Only proteins present in all 6 replicates of each pulldown were considered for this analysis. Finally, volcano plots for publication were generated using the VolcaNoseR web app ^32^.

### Sample Preparation for proteomics analysis of cell lines

Cell pellets of 30 M cells were lysed in lysis buffer (100 mM HEPES pH 7.6, 150 mM KCl, 1 mM MgCl2) by passing the cell suspension through a 27G needle 10 times using a 1 ml syringe. Subsequently, the cell lysate was incubated on ice for 20 minutes. The suspension was centrifuged for 4 minutes at 16000xg at 4°C and the supernatant was taken aside. Then the cell pellet was resuspended in lysis buffer and passed through a 27G neddle for 10 times. After 10 minutes incubation on ice, the suspension was centrifuged for 4 minutes at 16000xg at 4°C and the supernatant was mixed with the supernatant obtained from the first centrifugation. Lysate corresponding to 100 ug protein was subjected to digestion. After 5 minutes denaturation at 95°C, sodium deoxycholate (DOC) (Sigma-Aldrich) was added to a concentration of 5%. Proteins were reduced for 30 minutes at 37°C with 5 mM TCEP (Sigma-Aldrich) and then directly alkylated in the presence of 20 mM iodoacetamide (Sigma-Aldrich) for 40 minutes at room-temperature in the dark. Next, proteins were pre-digested for 4 hours at 37°C with 1 ug Lys-C (Fujifilm Wako Chemicals) and then diluted 1:5 in freshly prepared 0.1 M ammonium bicarbonate buffer to bring the concentration of DOC to 1%. Sequencing-grade Trypsin (Promega) was added at an enzyme:substrate ratio of 1:100 and incubated overnight at 37°C. At the next day, the tryptic digest was stopped and DOC was precipitated by the addition of formic acid. To remove DOC precipitates, the digest was centrifuged at 20000xg for 10 minutes and the supernatant was transferred to a fresh tube. This procedure was repeated twice. The cleared digest was loaded onto a 50 mg SepPak C18 column (Waters) which was primed with 100% methanol (Sigma-Aldrich), 80% ACN (Sigma-Aldrich) + 0.1% FA and equilibrated with 1% ACN + 0.1% FA. The flow-through was loaded one more time and then the column was washed three times with 1% ACN + 0.1% FA. Lastly, peptides were eluted in 35% ACN + 0.1% and dried in a SpeedVac for 4 hours at 45°C. For MS measurement peptides were reconstituted in 3% ACN + 0.1% FA.

### Liquid chromatography and mass spectrometry for proteomics analysis of cell lines

Peptides were analyzed in a data-independent acquisition mode (DIA) with an Exploris 480 (Thermo Scientific) mass spectrometer connected to an EASY-nLC (Thermo scientific) liquid chromatography system operating in nano-flow. Peptides were separated on a 30 cm fused silica column with 75 µm inner diameter packed in-house with 1.9 µm C18 beads (Dr. Maisch Reprosil-Pur 120). Peptides were separated along a 2 hours non-linear gradient constituting of a mixture of buffer A (3% ACN + 0.1% FA) and buffer B (90% ACN + 0.1% FA) at a flow rate of 250 nl/min. The DIA method acquired MS1 spectra with a scan range of 350-1650 m/z at a resolution of 120000 followed by 40 variable MS2 DIA windows with 0.5 m/z overlap at a resolution of 30000 and a normalized AGC Target of 3000%.

### Quantitative proteomics analysis for proteomics analysis of cell lines

DIA-MS runs were analyzed with Spectronaut 16 (Biognosys AG) in direct DIA+ search mode. The spectra were searched for Trypsin/P specific peptides with a length between 7-52 amino acids, allowing for two missed cleavages and setting carbamidomethyl(C) as fixed, and oxidation(M) and acetyl(protein, N-term) as variable modifications. Identifications were FDR controlled at 1% on precursor, peptide and protein level. The Uniprot release from 15.10.2020 of the *Homo sapiens* proteome was used as a reference. To search for peptides corresponding to PIPPI the fasta sequence of PIPPI was manually added to the reference proteome. The report table in the MSstats format was exported from Spectronaut 16 and further analyzed with MSstats (version 4.6.5) in R. Data from Spectronaut was filtered in MSstats using a q-value cutoff of 1% and removing proteins with only one feature. The data was normalized in MSstats using the “equalizeMedians” method and only the top 3 features were used to build quantities. For differential abundance testing, MSstats fits a linear mixed effects model and applies the Benjamini-Hochberg correction to account for multiple testing. Volcano plots were plotted in R using the ggplot package and. For the volcano plot, the significance thresholds were set to log2(FC) > 1 and and adjusted p-value < 0.05. The principal component analysis was done with base R using the MSstats protein quantities as input. The PCA plot was generated with ggplot in R.

## Data availability

Mass spectrometry data was uploaded to ProteomeXchange via PRIDE (PXD057806 and PXD058567). Reviewers log in using, pass `xFU2zZg4osEv` and, pass `gKsGvxyP7JB2`.

