## Supplementary Information for "Pooled overexpression screening identifies PIPPI as a novel microprotein involved in the ER stress response"

**Supplementary Figures 1-5**

**Supplementary Data**

Location and genomic sequence of PIPPI sORFs

**Supplementary Tables 1-4**

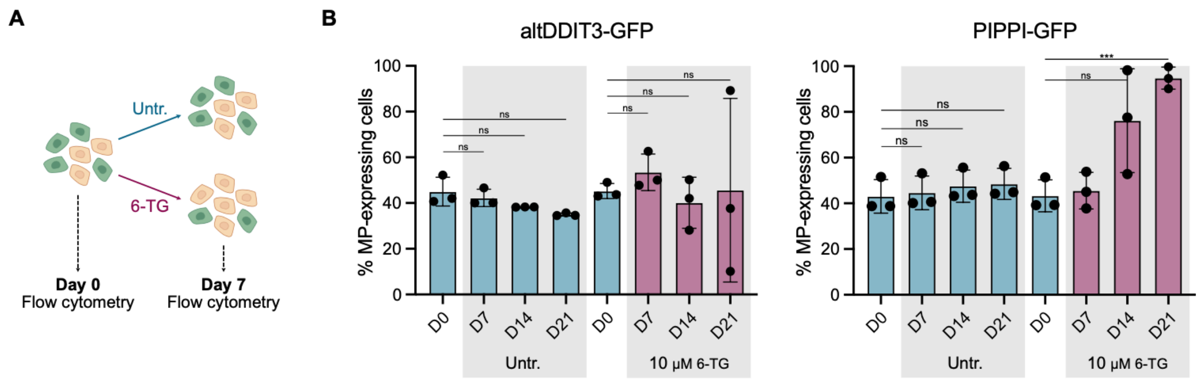


**Supplementary Figure 1**

A) Schema illustrating the design of the growth competition assays performed to validate the results of the pooled overexpression screen. In short, microprotein-expressing cells (beige cells) are mixed 1:1 with their parental counterparts expressing a fluorescent marker (green cells). The mixed populations are either cultured untreated for 7 days or left recover for the same period following a 24 hours 6-TG treatment. At day 0 and day 7 of the assay, cells are collected, and the amount of GFP-positive and GFP-negative cells present in each sample is assessed by flow cytometry. B) Results of the growth competition assays comparing the growth of altDDIT3-GFP or PIPPI-GFP to A375 parental cells. Where indicated, cells were continuously treated with 10 mM 6-TG, which was renewed every 2-3 days until the end of the assay. Height of the bars represents the fraction of microprotein-expressing cells present in the total cell population. Values were averaged based on 3 independent biological replicates (black circles). p-values were calculated using unpaired Student's t-test (ns = $p > 0.05$, *** = $p < 0.001$).

Figure S2

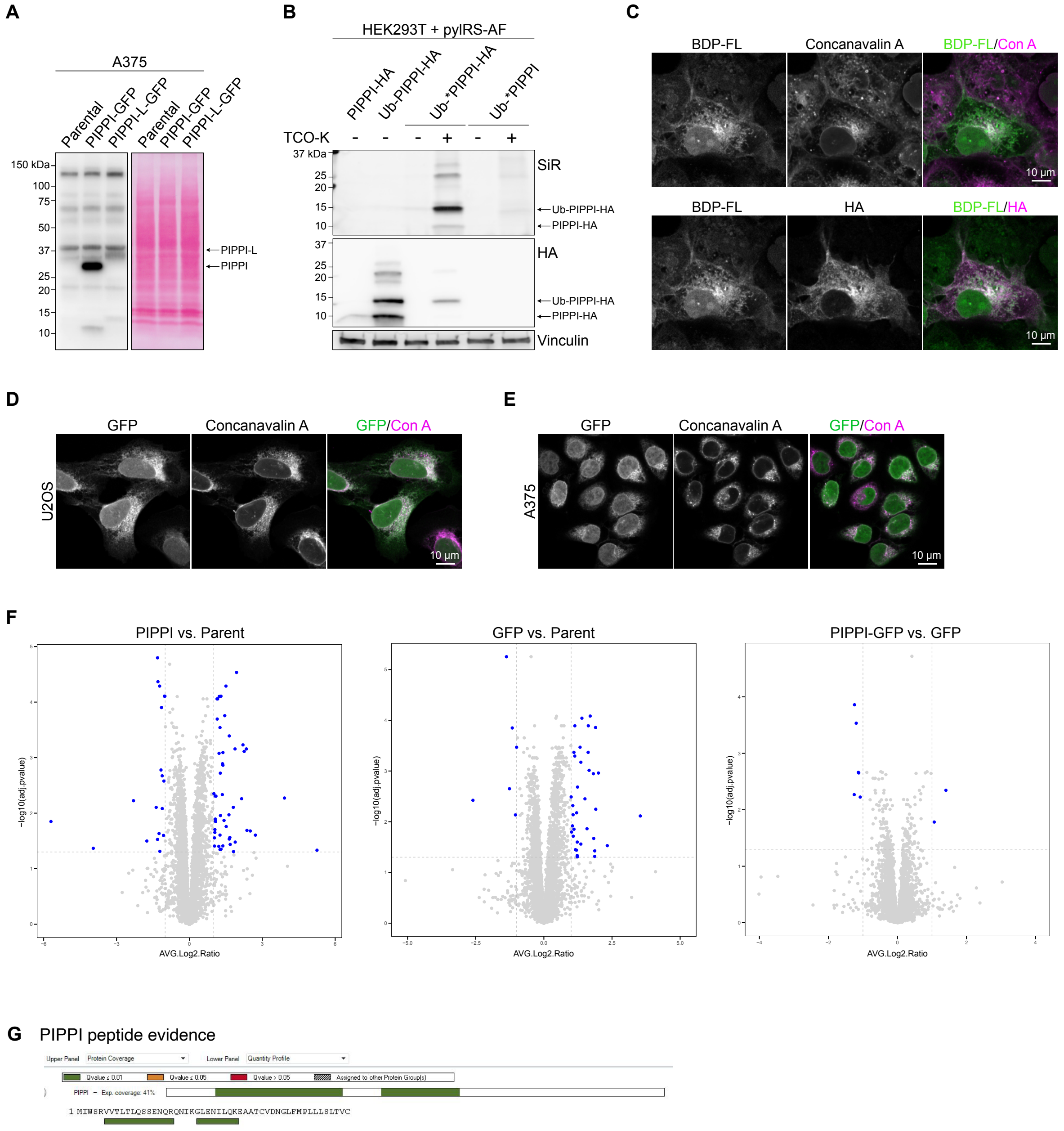


**Supplementary Figure 2**

A) Immunoblots showing the levels of PIPPI-GFP and PIPPI-L-GFP overexpression in A375 cells. B) Various PIPPI-HA constructs were transiently expressed in HEK293T cells. Where indicated, the pyrrolysine tRNA synthetase (pylRS) and the non-canonical amino acid TCO*K were provided to allow suppression of the amber stop codon present in some of the constructs. Lysates were labelled using SiR tetrazine and analysed by SDS-PAGE followed by immunoblotting. C) Images presented in Figure 2C are overlapped to highlight the extent of colocalization existing between the BDP-FL and HA signals, as well as the BFP-FL and Concanavalin A (Con A) signals. D-E) Confocal images of U2OS (D) and A375 (E) cells stably expressing a PIPPI-GFP transgene. Concavalin A (Con A) staining was included to visualize the endoplasmic reticulum. F) Volcano plots presenting the pairwise comparisons of overexpression and the control cell lines. Thresholds are set at Log2(FC) = 1 and adjusted p-value = 0.05 (dashed lines). G) Peptide evidence for PIPPI.

### Figure S3

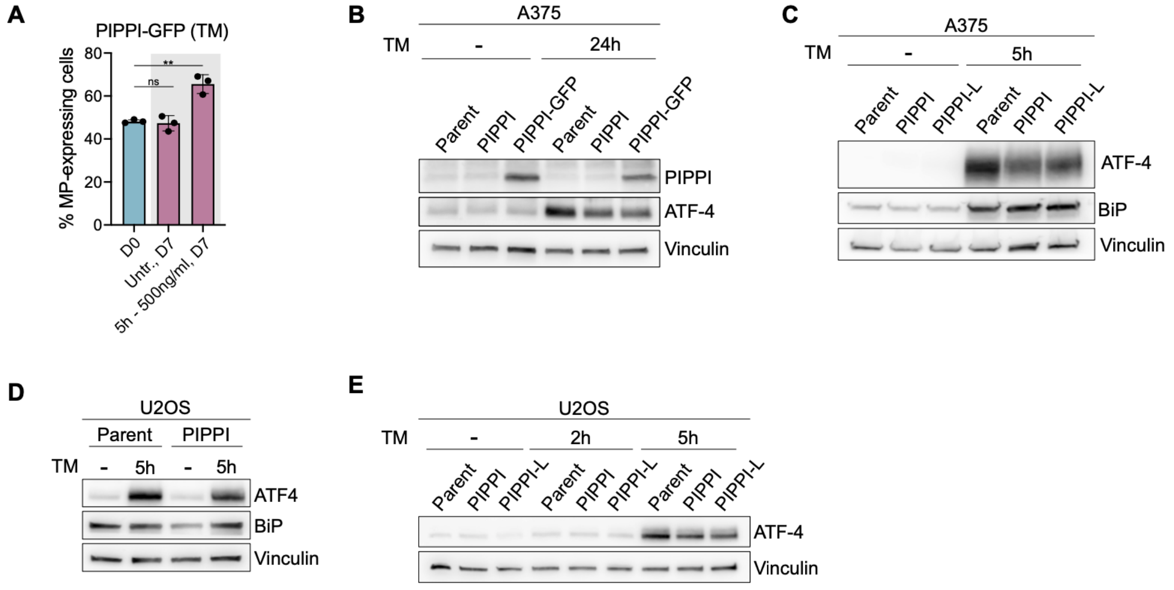


**Supplementary Figure 3**

A) Bar chart presenting the results of the growth competition assays performed to assess the ability of PIPPI-GFP-expressing cells to overcome tunicamycin treatment. Height of the bars represents the fraction of PIPPI-GFP-expressing cells present in the total cell population. Values were averaged based on 3 independent biological replicates (black circles). p-values were calculated using unpaired Student's t-test (ns = $p > 0.05$; ** = $p < 0.01$). B) Parental, PIPPI- and PIPPI-GFP-expressing A375 cells were treated with 100 ng/ml Tunicamycin for 24 hours. Lysates were then analysed by immunoblotting. C) Parental, PIPPI- and PIPPI-L-expressing A375 cells were treated for 5h with 500 ng/ml Tunicamycin. Induction of the UPR was assessed by immunoblotting. D) Parental and PIPPI-expressing U2OS cells were treated with 500 ng/ml Tunicamycin for 5 hours. After lysis, the levels of ATF4, BiP and vinculin were assessed by immunoblotting. E) Parental, PIPPI- and PIPPI-L-expressing U2OS cells were treated with 200 ng/ml Tunicamycin for either 2 or 5 hours. The extent of ATF4 induction was then assessed by immunoblotting.

Figure S4

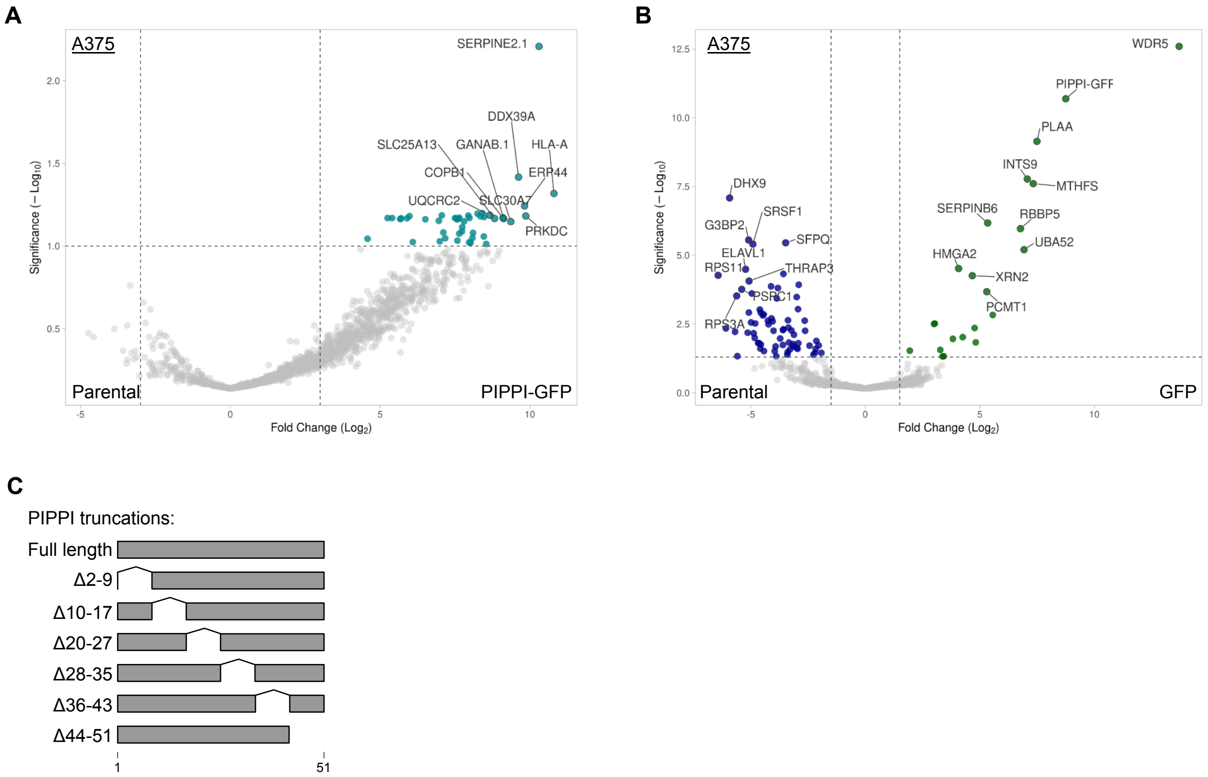


**Supplementary Figure 4**

A) Volcano plot of co-immunoprecipitation LC-MS/MS (co-IP/MS) experiments performed with parental and PIPPI-GFP-expressing A375 cells. Experiment was conducted in two biological replicates, which were each analyzed three times by mass-spectrometry. Thresholds are set at Log2FC = 3 and DEP adjusted p-value = 0.1 (dashed lines). Proteins enriched in PIPPI-GFP lysates are clustering in the right half of the plot and the top ten proteins (assessed by Manhattan distance) are labelled with their names. B) Volcano plot of co-immunoprecipitation LC-MS/MS (co-IP/MS) experiments performed with parental and GFP-expressing A375 cells. Experiment was conducted in two biological replicates, which were each analyzed three times by mass-spectrometry. Thresholds are set at Log2FC = 1. and DEP adjusted p-value = 0.1 (dashed lines). Proteins significantly enriched in parental lysates are labeled in blue, whereas proteins enriched in GFP lysates are labeled in green. In both cases, the top ten proteins (assessed by Manhattan distance) are labelled with their names. C) Schema depicting the different PIPPI-GFP truncations that were tested in Figure 4 E-F.

Figure S5

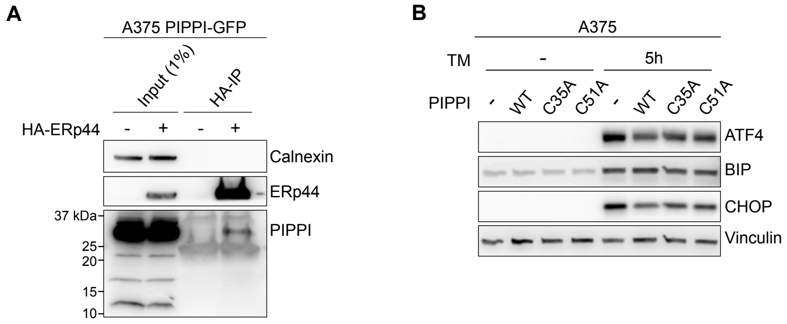


**Supplementary Figure 5**

A) Lysates from A375 cells either expressing PIPPI-GFP or co-expressing PIPPI-GFP and HA-ERp44 were subjected to immunoprecipitation using HA-beads and analysed by immunoblotting with the antibodies indicated on the right. B) Parental, wild type PIPPI-GFP-, C35A PIPPI-GFP-, and C51A PIPPI-GFP-expressing A375 cells were treated with 500 ng/ml Tunicamycin for 5 hours. After lysis, the level of different proteins involved in the ER stress response were assessed by immunoblotting.

### Supplementary Data – Location and genomic sequences of PIPPI sORFs

#### Ensembl release 110 - July 2023

| | |
|---|---|
| **Exons** | All exons |
| **HSP** | Location of selected alignment |
| **Markup** | loaded |

**>chromosome:GRCh38:16:15104848:15105597:-1**
15105597 CCTGGGAGTGAAAAGAAATATTACAGCCATGCCTAACTGACTTCTTGAGGTAAGATTGTT 15105538
15105537 CTGTCAGAAAACCCTCTCCCAGTTCCCCTGCAGCTCTTCAGGAATCCACATCTCTCCAGA 15105478
15105477 GCTCTTTGTTCTCATGGGTGGCACCTCCAGAGTGAAGAAGATCCTTTGTCAAGAAGGGAA 15105418
15105417 ACAGAGGGGAAATGAGAGGGTCCTGCAGGCAGAGCTGGAATCAACTTCCACTCTGCCTCT 15105358
15105357 TGCAAGCTGTGTGACCCTGGGCACAATTTCTCCTTCCTCTGGAAACCTCTGTTTTCTTAG 15105298
15105297 **ATTTGGAGCAGGGTGGTCACACTGACCTTGCAGAGTTCTGAGAATCAGAGACAGAACATA** 15105238
15105237 **AAGGGCCTGGAAAACATTCTCCAAAAAGAAGCTGCAACATGTGTGGACAATGGGCTTTTC** 15105178
15105177 **ATGCCTCTCTTACTGTCTCTTACTGTCTGT**TGACCTGGTGCAAGAAACATGCTCTGGTGA 15105118
15105117 TGGCTGTGAGGGAGGAATGAGGATAGACATAGACACTCCTGTGTCTCAAACATGCTTCTT 15105058
15105057 TATTACTCTGTTATGACTCTGTCTTCCCTGGGGCAGGACCCCAGCCTGCCTACATTTGCA 15104998
15104997 GACAGACACAGTGGCATGTGGAGACAACAGTGTGTCCCAGTGACTTTTCTTTACCCCCCA 15104938
15104937 GCTGTCGGCAGTACTCAGTGGAAGGGTGATATGACACTGATACTGCTATTTTGAAACCTG 15104878
15104877 GAGGATGGAAAGGTGCAAAAATCTATCACC 15104848

**>chromosome:GRCh38:16:11927972:11928721:-1**
11928721 CCTGGGAGTGAAAAGAAATATTACAGCCATGCCTAACTGACTTCTTGAGGTGAGATTGTT 11928662
11928661 CTGTCAGAAAACCCTCTCCCAGTTCCCCTGCAGCTCTTCAGGAATCCACATCTCTCCAGA 11928602
11928601 GCTCTTTGTTCTCATGGGTGGCACCTCCAGAGTGAAGAAGATCCTTTGTCAAGAAGGGAA 11928542
11928541 ACAGAGGGGAAATGAGAGGGTCCTGCAGGCAGAGCTGGAATCAACTTCCACTCTGCCTCT 11928482
11928481 TGCAAGCTGTGTGACCCTGGGCACAATTTCTCCTTCCTCTGGAAACCTCTGTTTTCTTAG 11928422
11928421 **ATTTGGAGCAGGGTGGTCACACTGACCTTGCAGAGTTCTGAGAATCAGAGACAGAACATA** 11928362
11928361 **AAAGGCCTGGAAAACATTCTCCAAAAAGAAGCTGCAACATGTGTGGACAATGGGCTTTTC** 11928302
11928301 **ATGCCTCTCTTACTGTCTCTTACTGTCTGT**TGACCTGGTGCAAGAAACATGCTCTGGTGA 11928242
11928241 TGGCTGTGAGGGAGGAATGAGGATAGACATAGACACTCCTGTGTCTCAAACATGCTTCTT 11928182
11928181 TATTACTCTGTTATGACTCTGTCTTCCCTGGGGCAGGACCCCAGCCTGCCTACATTTGCA 11928122
11928121 GACAGACACAGTGGCATGTGGAGACAACAGTGTGTCCCAGTGACTTTTCTTTACCCTCCA 11928062
11928061 GCTGTTGGCAGTACTCAGTGGAAGGGTGATATTATGACACTGATACTGCTATTTTGAAAC 11928002
11928001 CTGGAGGATGGAAAGGTGCAAAAATCTATC 11927972

**>chromosome:GRCh38:16:22532784:22533533:1**
22532784 CCTGGGAGTGAAAAGAAATATTACAGCCATGCCTAAGTGACTTCTTGAGGTGAGATTGTT 22532843
22532844 CTGTCAGAAAACCCTCTCCCAGTTCCCCTGCAGCTCTTCAGGAATCCACATCTCTCCAGA 22532903
22532904 GCTCTTTGTTCTCATGGGTGGCACCTCCAGAGTGAAGAAGATCCTTTGTCAAGAAGGGAA 22532963
22532964 ACAGAAGGGAAATGAGAGGGTCCTGCAGGCAGAGCTGGAATCAACTTCCACTCTGCCTCT 22533023
22533024 TGCAAGCTGTGTGACCCTGGGCACAATTTCTCCTTCCTCTGGAAACCTCTGTTTTCTTAG 22533083
22533084 **ATTTGGAGCAGGGTGGTCACACTGACCTTGCAGAGTTCTGAGAATCAGAGACAGAACATA** 22533143
22533144 **AAAGGCCTGGAAAACATTCTCCAAAAAGAAGCTGCAACATGTGTGGACAGTGGGCTTTTC** 22533203
22533204 **ATGCCTCTCTTACTGTCTCTTACTGTCTGT**TGACCTGGTGCAAGAAACATGCTCTGGTGA 22533263
22533264 TGGCTGTGAGGGAGGAATGAGGATAGACATAGACACTCCTGTGTCTCAAACATGCTTCTT 22533323
22533324 TATTACTCTGTTATGACTCTGTCTTCCCTGGGGCAGGACCCCAGCCTGCCTACATTTGCA 22533383
22533384 GACAGACACAGTGGCATGTGGAGACAACAGTGTGTCCCAATGACTTTCCTTTACCCTCCA 22533443
22533444 GCTGTCGGCAGTACTCAGTGGAAGGGTGATATTATGACACTGATACTGCTATTTTGAAAC 22533503
22533504 CTGGAGGATGGAAAGGTGCAAAAATCTATC 22533533

**>chromosome:GRCh38:16:28771311:28772060:1**
28771311 CCTGGGAGTGAAAAGAAATATTACAGCCATGCCTAAGTGACTTCTTGAGGTGAGATTGTT 28771370
28771371 CTGTCAGAAAACCCTCTCCCAGTTCCCCTGCAGCTCTTCAGGAATCCACATCTCTCCAGA 28771430
28771431 GCTCTTTGTTCTCATGGGTGGCACCTCCAGAGTGAAGAAGATCCTTTGTCAAGAAGGGAA 28771490
28771491 ACAGAGGGGAAATGAGAGGGTCCTTCAGGCAGAGCTGGAATCAACTTCCACTCTGCCTCT 28771550
28771551 TGCAAGCTGTGTGACCCTGGGCACAATTTCTCCTTCCTCTGGAAACCTCTGTTTTCTTAG 28771610
28771611 **ATTTGGAGCAGGGTGGTCACACTGACCTTGCAGAGTTCTGAGAGTCAGAGACAGAACATA** 28771670
28771671 **AAAGGCCTGGAAAACATTCTCCAAAAAGAAGCTGCAACATGTGTGGACAATGGGCTTTTC** 28771730

28771731 **ATGCCTCTCTTACTGTCTCTTACTGTCTGT**TGACCTGGTGCAAGAAACATGCTCTGGTGA 28771790
28771791 TGGCTGTGAGGGAGGAATGAGGATAGACATAGACACTCCTGTGTCTCAAACATGCTTCTT 28771850
28771851 TATTACTCTGTTATGACTCTGTCTTCCCTGGGGCAGGACCCCAGCCTGCCTACATTTGCA 28771910
28771911 GACAGACACAGTGGCATGTGGAGACAACAGTGTGTCCCAATGACTTTCCTTTACCCTCCA 28771970
28771971 GCTGTCGGCAGTACTCAGTGGAAGGGTGATATTATGACACTGATACTGCTATTTTGAAAC 28772030
28772031 CTGGAGGATGGAAAGGTGCAAAAATCTATC 28772060

**>chromosome:GRCh38:16:29051230:29051979:1**
29051230 GGAGTGAAAAGAAATATTACAGCCATGCATGCCTAAGTGACTTCTTGAGGTGAGATTGTT 29051289
29051290 CTGTCAGAAAACCCTCTCCCAGTTCCCCTGCAGCTCTTCAGGAATCCACATCTCTCCAGA 29051349
29051350 GCTCTTTGTTCTCATGGGTGGCACCTCCAGAGTGAAGAAGATCCTTTGTCAAGAAGGGAA 29051409
29051410 ACAGAGGGGAAATGAGAGGGTCCTGCAGGCAGAGCTGGAATCAACTTCCACTCTGCCTCT 29051469
29051470 TGCAAGCTGTGTGACCCTGGGCACAATTTCTCCTTCCTCTGGAAACCTCTGTTTTCTTAG 29051529
29051530 **ATTTGGAGCAGGGTGGTCACACTGACCTTGCAGAGTTCTGAGAGTCAGAGACAGAACATA** 29051589
29051590 **AAAGGCCTGGAAAACATTCTCCAAAAAGAAGCTGCAACATGTGTGGACAATGGGCTTTTC** 29051649
29051650 **ATGCCTCTCTTACTGTCTCTTACTGTCTGT**TGACCTGGTGCAAGAAACATGCTCTGGTGA 29051709
29051710 TGGCTGTGAGGGAGGAATGAGGATAGACATAGACACTCCTGTGTCTCAAACATGCTTCTT 29051769
29051770 TATTACTCTGTTATGACTCTGTCTTCCCTGGGGCAGGACCCCAGCCTGCCTACATTTGCA 29051829
29051830 GACAGACACAGTGGCATGTGGAGACAACAGTGTGTCCCAATGACTTTTCTTTACCCCCCA 29051889
29051890 GCTGTCGGCAGTACTCAGTGGAAGGGTGATATTATGACACTGATACTGCTATTTTGAAAC 29051949
29051950 CTGGAGGATGGAAAGGTGCAAAAATCTATC 29051979

**>chromosome:GRCh38:16:29384382:29385131:-1**
29385131 CCTGGGAGTGAAAAGAAATATTACAGCCATGCCTAAGTGACTTCTTGAGGTGAGATTGTT 29385072
29385071 CTGTCAGAAAACCCTCTCCCAGTTCCCCTGCAGCTCTTCAGGAATCCACATCTCTCCAGA 29385012
29385011 GCTCTTTGTTCTCATGGGTGGCACCTCCAGAGTGAAGAAGATCCTTTGTCAAGAAGGGAA 29384952
29384951 ACAGAGGGGAAATGAGAGGGTCCTGCAGGCAGAGCTGGAATCAACTTCCACTCTGCCTCT 29384892
29384891 TGCAAGCTGTGTGACCCTGGGCACAATTTCTCCTTCCTCTGGAAACCTCTGTTTTCTTAG 29384832
29384831 **ATTTGGAGCAGGGTGGTCACACTGACCTTGCAGAGTTCTGAGAATCAGAGACAGAACATA** 29384772
29384771 **AAAGGCCTGGAAAACATTCTCCAAAAAGAAGCTGCAACATGTGTGGACAGTGGGCTTTTC** 29384712
29384711 **ATGCCTCTCTTACTGTCTCTTACTGTCTGT**TGACCTGGTGCAAGAAACATGCTCTGGTGA 29384652
29384651 TGGCTGTGAGGGAGGAATGAGGATAGACATAGACACTCCTGTGTCTCAAACATGCTTCTT 29384592
29384591 TATTACTCTGTTATGACTCTGTCTTCCCTGGGGCAGGACCCCAGCCTGCCTACATTTGCA 29384532
29384531 GACAGACACAGTGGCATGTGGAGACAACAGTGTGTCCCAATGACTTTCCTTTACCCTCCA 29384472
29384471 GCTGTCGGCAGTACTCAGTGGAAGGGTGATATTATGACACTGATACTGCTATTTTGAAAC 29384412
29384411 CTGGAGGATGGAAAGGTGCAAAAATCTATC 29384382

**>chromosome:GRCh38:16:28343281:28344030:-1**
28344030 CCTGGGAGTGAAAAGAAATATTACAGCCATGCCTAAGTGACTTCTTGAGGTGAGATTGTT 28343971
28343970 CTGTCAGAAAACCCTCTCCCAGTTCCCCTGCAGCTCTTCAGGAATCCACATCTCTCCAGA 28343911
28343910 GCTCTTTGTTCTCATGGGTGGCACCTCCAGAGTGAAGAAGATCCTTTGTCAAGAAGGGAA 28343851
28343850 ACAGAGGGGAAATGAGAGGGTCCTGCAGGCAGAGCTGGAATCAACTTCCACTCTGCCTCT 28343791
28343790 TGCAAGCTGTGTGACCCTGGGCACAATTTCTCCTTCCTCTGGAAACCTCTGTTTTCTTAG 28343731
28343730 **ATTTGGAGCAGGGTGGTCACACTGACCTTGCAGAGTTCTGAGAGTCAGAGACAGAACATA** 28343671
28343670 **AAAGGCCTGGAAAACATTCTCCAAAAAGAAGCTGCAACATGTGTGGACAATGGGCTTTTC** 28343611
28343610 **ATGCCTCTCTTACTGTCTCTTACTGTCTGT**TGACCTGGTGCAAGAAACATGCTCTGGTGA 28343551
28343550 TGGCTGTGAGGGAGGAATGAGGATAGACATAGACACTCCTGTGTCTCAAACATGCTTCTT 28343491
28343490 TATTACTCTGTTATGACTCTGTCTTCCCTGGGGCAGGACCCCAGCCTGCCTACATTTGCA 28343431
28343430 GACAGACACAGTGGCATGTGGAGACAACAGTGTGTCCCAATGACTTTTCTTTACCCCCCA 28343371
28343370 GCTGTCGGCAGTACTCAGTGGAAGGGTGATATTATGACACTGATACTGCTATTTTGAAAC 28343311
28343310 CTGGAGGATGGAAAGGTGCAAAAATCTATC 28343281

**>chromosome:GRCh38:16:21837837:21838586:-1**
21838586 CCTGGGAGTGAAAAGAAATATTACAGCCATGCCTAAGTGACTTCTTGAGGTGAGATTGTT 21838527
21838526 CTGTCAGAAAACCCTCTCCCAGTTCCCCTGCAGCTCTTCAGGAATCCACATCTCTCCAGA 21838467
21838466 GCTCTTTGTTCTCATGGGTGGCACCTCCAGAGTGAAGAAGATCCTTTGTCAAGAAGGGAA 21838407
21838406 ACAGAGGGGAAATGAGAGGGTCCTGCAGGCAGAGCTGGAATCAACTTCCACTCTGCCTCT 21838347
21838346 TGCAAGCTGTGTGACCCTGGGCACAATTTCTCCTTCCTCTGGAAACCTCTGTTTTCTTAG 21838287
21838286 **ATTTGGAGCAGGGTGGTCACACTGACCTTGCAGAGTTCTGAGAATCAGAGACAGAACATA** 21838227
21838226 **AAAGGCCTGGAAAACATTCTCCAAAAAGAAGCTGCAACATGTGTGGACAGTGGGCTTTTC** 21838167
21838166 **ATGCCTCTCTTACTGTCTCTTACTGTCTGT**TGACCTGGTGCAAGAAACATGCTCTGGTGA 21838107
21838106 TGGCTGTGAGGGAGGAATGAGGATAGACATAGACACTCCTGTGTCTCAAACATGCTTCTT 21838047

21838046 TATTACTCTGTTATGACTCTGTCTTCCCTGGGGCAGGACCCCAGCCTGCCTACATTTGCA 21837987
21837986 GACAGACACAGTGGCATGTGGAGACAACAGTGTGTCCCAATGACTTTCCTTTACCCTCCA 21837927
21837926 GCTGTTGGCAGTACTCAGTGGAAGGGTGATATTATGACACTGATACTTCTATTTTGAAAC 21837867
21837866 CTGGAGGATGGAAAGGTGCAAAAATCTATC 21837837

**>chromosome:GRCh38:16:21405386:21406135:-1**
21406135 CCTGGGAGTGAAAAGAAATATTACAGCCATGCCTAAGTGACTTCTTGAGGTGAGATTGTT 21406076
21406075 CTGTCAGAAAACCCTCTCCCAGTTCCCCTGCAGCTCTTCAGGAATCCACATCTCTCCAGA 21406016
21406015 GCTCTTTGTTCTCATGGGTGGCACCTCCAGAGTGAAGAAGATCCTTTGTCAAGAAGGGAA 21405956
21405955 ACAGAGGGGAAATGAGAGGGTCCTGCAGGCAGAGCTGGAATCAACTTCCACTCTGCCTCT 21405896
21405895 TGCAAGCTGTGTGACCCTGGGCACAATTTCTCCTTCCTCTGGAAACCTCTGTTTTCTTAG 21405836
21405835 **ATTTGGAGCAGGGTGGTCACACTGACCTTGCAGAGTTCTGAGAATCAGAGACAGAACATA** 21405776
21405775 **AAAGGCCTGGAAAACATTCTCCAAAAAGAAGCTGCAACATGTGTGGACAGTGGGCTTTTC** 21405716
21405715 **ATGCCTCTCTTACTGTCTCTTACTGTCTGT**TGACCTGGTGCAAGAAACATGCTCTGGTGA 21405656
21405655 TGGCTGTGAGGGAGGAATGAGGATAGACATAGACACTCCTGTGTCTCAAACATGCTTCTT 21405596
21405595 TATTACTCTGTTATGACTCTGTCTTCCCTGGGGCAGGACCCCAGCCTGCCTACATTTGCA 21405536
21405535 GACAGACACAGTGGCATGTGGAGACAACAGTGTGTCCCAATGACTTTCCTTTACCCTCCA 21405476
21405475 GCTGTCGGCAGTACTCAGTGGAAGGGTGATATTATGACACTGATACTGCTATTTTGAAAC 21405416
21405415 CTGGAGGATGGAAAGGTGCAAAAATCTATC 21405386

**>scaffold:GRCh38:HG926_PATCH:1266441:1267190:1**
1266441 CCTGGGAGTGAAAAGAAATATTACAGCCATGCCTAAGTGACTTCTTGAGGTGAGATTGTT 1266500
1266501 CTGTCAGAAAACCCTCTCCCAGTTCCCCTGCAGCTCTTCAGGAATCCACATCTCTCCAGA 1266560
1266561 GCTCTTTGTTCTCATGGGTGGCACCTCCAGAGTGAAGAAGATCCTTTGTCAAGAAGGGAA 1266620
1266621 ACAGAGGGGAAATGAGAGGGTCCTGCAGGCAGAGCTGGAATCAACTTCCACTCTGCCTCT 1266680
1266681 TGCAAGCTGTGTGACCCTGGGCACAATTTCTCCTTCCTCTGGAAACCTCTGTTTTCTTAG 1266740
1266741 **ATTTGGAGCAGGGTGGTCACACTGACCTTGCAGAGTTCTGAGAATCAGAGACAGAACATA** 1266800
1266801 **AAAGGCCTGGAAAACATTCTCCAAAAAGAAGCTGCAACATGTGTGGACAGTGGGCTTTTC** 1266860
1266861 **ATGCCTCTCTTACTGTCTCTTACTGTCTGT**TGACCTGGTGCAAGAAACATGCTCTGGTGA 1266920
1266921 TGGCTGTGAGGGAGGAATGAGGATAGACATAGACACTCCTGTGTCTCAAACATGCTTCTT 1266980
1266981 TATTACTCTGTTATGACTCTGTCTTCCCTGGGGCAGGACCCCAGCCTGCCTACATTTGCA 1267040
1267041 GACAGACACAGTGGCATGTGGAGACAACAGTGTGTCCCAATGACTTTCCTTTACCCTCCA 1267100
1267101 GCTGTCGGCAGTACTCAGTGGAAGGGTGATATTATGACACTGATACTTCTATTTTGAAAC 1267160
1267161 CTGGAGGATGGAAAGGTGCAAAAATCTATC 1267190

**>scaffold:GRCh38:HG926_PATCH:571504:572253:-1**
572253 CCTGGGAGTGAAAAGAAATATTACAGCCATGCCTAAGTGACTTCTTGAGGTGAGATTGTT 572194
572193 CTGTCAGAAAACCCTCTCCCAGTTCCCCTGCAGCTCTTCAGGAATCCACATCTCTCCAGA 572134
572133 GCTCTTTGTTCTCATGGGTGGCACCTCCAGAGTGAAGAAGATCCTTTGTCAAGAAGGGAA 572074
572073 ACAGAGGGGAAATGAGAGGGTCCTGCAGGCAGAGCTGGAATCAACTTCCACTCTGCCTCT 572014
572013 TGCAAGCTGTGTGACCCTGGGCACAATTTCTCCTTCCTCTGGAAACCTCTGTTTTCTTAG 571954
571953 **ATTTGGAGCAGGGTGGTCACACTGACCTTGCAGAGTTCTGAGAATCAGAGACAGAACATA** 571894
571893 **AAAGGCCTGGAAAACATTCTCCAAAAAGAAGCTGCAACATGTGTGGACAGTGGGCTTTTC** 571834
571833 **ATGCCTCTCTTACTGTCTCTTACTGTCTGT**TGACCTGGTGCAAGAAACATGCTCTGGTGA 571774
571773 TGGCTGTGAGGGAGGAATGAGGATAGACATAGACACTCCTGTGTCTCAAACATGCTTCTT 571714
571713 TATTACTCTGTTATGACTCTGTCTTCCCTGGGGCAGGACCCCAGCCTGCCTACATTTGCA 571654
571653 GACAGACACAGTGGCATGTGGAGACAACAGTGTGTCCCAATGACTTTCCTTTACCCTCCA 571594
571593 GCTGTCGGCAGTACTCAGTGGAAGGGTGATATTATGACACTGATACTGCTATTTTGAAAC 571534
571533 CTGGAGGATGGAAAGGTGCAAAAATCTATC 571504

**>scaffold:GRCh38:HG926_PATCH:228351:229100:-1**
229100 CCTGGGAGTGAAAAGAAATATTACAGCCATGCCTAAGTGACTTCTTGAGGTGAGATTGTT 229041
229040 CTGTCAGAAAACCCTCTCCCAGTTCCCCTGCAGCTCTTCAGGAATCCACATCTCTCCAGA 228981
228980 GCTCTTTGTTCTCATGGGTGGCACCTCCAGAGTGAAGAAGATCCTTTGTCAAGAAGGGAA 228921
228920 ACAGAGGGGAAATGAGAGGGTCCTGCAGGCAGAGCTGGAATCAACTTCCACTCTGCCTCT 228861
228860 TGCAAGCTGTGTGACCCTGGGCACAATTTCTCCTTCCTCTGGAAACCTCTGTTTTCTTAG 228801
228800 **ATTTGGAGCAGGGTGGTCACACTGACCTTGCAGAGTTCTGAGAATCAGAGACAGAACATA** 228741
228740 **AAAGGCCTGGAAAACATTCTCCAAAAAGAAGCTGCAACATGTGTGGACAGTGGGCTTTTC** 228681
228680 **ATGCCTCTCTTACTGTCTCTTACTGTCTGT**TGACCTGGTGCAAGAAACATGCTCTGGTGA 228621
228620 TGGCTGTGAGGGAGGAATGAGGATAGACATAGACACTCCTGTGTCTCAAACATGCTTCTT 228561
228560 TATTACTCTGTTATGACTCTGTCTTCCCTGGGGCAGGACCCCAGCCTGCCTACATTTGCA 228501
228500 GACAGACACAGTGGCATGTGGAGACAACAGTGTGTCCCAATGACTTTCCTTTACCCTCCA 228441
228440 GCTGTCGGCAGTACTCAGTGGAAGGGTGATATTATGACACTGATACTGCTATTTTGAAAC 228381
228380 CTGGAGGATGGAAAGGTGCAAAAATCTATC 228351

**>scaffold:GRCh38:HG926_PATCH:1698628:1699377:1**
1698628 CCTGGGAGTGAAAAGAAATATTACAGCCATGCCTAAGTGACTTCTTGAGGTGAGATTGTT 1698687
1698688 CTGTCAGAAAACCCTCTCCCAGTTCCCCTGCAGCTCTTCAGGAATCCACATCTCTCCAGA 1698747
1698748 GCTCTTTGTTCTCATGGGTGGCACCTCCAGAGTGAAGAAGATCCTTTGTCAAGAAGGGAA 1698807
1698808 ACAGAGGGGAAATGAGAGGGTCCTGCAGGCAGAGCTGGAATCAACTTCCACTCTGCCTCT 1698867
1698868 TGCAAGCTGTGTGACCCTGGGCACAATTTCTCCTTCCTCTGGAAACCTCTGTTTTCTTAG 1698927
1698928 **ATTTGGAGCAGGGTGGTCACACTGACCTTGCAGAGTTCTGAGAATCAGAGACAGAACATA** 1698987
1698988 **AAAGGCCTGGAAAACATTCTCCAAAAAGAAGCTGCAACATGTGTGGACAGTGGGCTTTTC** 1699047
1699048 **ATGCCTCTCTTACTGTCTCTTACTGTCTGT**TGACCTGGTGCAAGAAACATGCTCTGGTGA 1699107
1699108 TGGCTGTGAGGGAGGAATGAGGATAGACATAGACACTCCTGTGTCTCAAACATGCTTCTT 1699167
1699168 TATTACTCTGTTATGACTCTGTCTTCCCTGGGGCAGGACCCCAGCCTGCCTACATTTGCA 1699227
1699228 GACAGACACAGTGGCATGTGGAGACAACAGTGTGTCCCAATGACTTTCCTTTACCCTCCA 1699287
1699288 GCTGTCGGCAGTACTCAGTGGAAGGGTGATATTATGACACTGATACTGCTATTTTGAAAC 1699347
1699348 CTGGAGGATGGAAAGGTGCAAAAATCTATC 1699377

**>chromosome:GRCh38:16:74390549:74391298:1**
74390549 CCTGGGAGTGAAAAGAAATATTACAGCCATGCCTAACTGACTTCTTGAGGTAAGATTGTT 74390608
74390609 CTGTCAGAAAACCCTCTCCCAGTTCCCCTGCAGCTCTTCAGGAACCCACATCTCTCCAGA 74390668
74390669 GCTCTTTGTTCTCATGGGTGGCACCTCCAGAGTGAAGAAGATCCTTTGTCAAGAAGGGAA 74390728
74390729 ACAGAGGGGAAATGAGAGGGTCCTGCAGGCAGAGCTGGAATCAACTTCCACTCTGCCTCT 74390788
74390789 TGCAAGCTGTGTGACCCTGGGCACAATTTCTCCTTCCTCTGGAAACCTCTGTTTTCTTAG 74390848
74390849 **ATTTGGAGCAGGGTGGTCACACTGACCTTGCAGAGTCCTGAGAATCAGAGACAGAACATA** 74390908
74390909 **AAAGGCCTGGAAAACATTCTCCAAAAAGAAGCTGCAACATGTGTGGACAATGGGCTTTTC** 74390968
74390969 **ATGCCTCTCTTACTGTCTCTTACTGTCTGT**TGACCTGGTGCAAGAAACATGCTCTGGTGA 74391028
74391029 TGGCTGTGAGGGAGGAATGAGGATAGACATAGACACTCCTGTGTCTCAAACATGCCTCTT 74391088
74391089 TATTACTCTGTTATGACTCTGTCTTCCCTGGGGCAGGACCCCAGCCTGCCTACATTTGCA 74391148
74391149 GACAGACACAGTGGCATGTGGAGACAACAGTGTGTCCCAATGACTTTTCTTTACTCCCCA 74391208
74391209 GCTGTCGGCAGTACTCAGTGGAAGGGTGATATCATGACACTGATACTGCTATTTTGAAAC 74391268
74391269 CTGGAGGATGGAAAGGTGCAAAAATCTATC 74391298

**>chromosome:GRCh38:16:14725041:14725787:1**
14725041 CCTGGGAGTGAAAAGAAATATTACAGCCATGCCTAACTGACTTCTTGAGGTAAGATTGTT 14725100
14725101 CTGTCAGAAAACCCTCTCCCAGTTCCCCTGCAGCTCTTCAGGAATCCACATCTCTCCAGA 14725160
14725161 GCTCTTTGTTCTCATGGGTGGCACCTCCAGAGTGAAGAAGATCCTTTGTCAAGAAGGGAA 14725220
14725221 ACAGAGGGGAAATGAGAGGGTCCTGCAGGCAGAGCTGGAATCAACTTCCACTCTGCCTCT 14725280
14725281 TGCAAGCTGTGTGACCCTGGGCACAATTTCTCCTTCCTCTGGAAACCTCTGTTTTCTTAG 14725340
14725341 **ATTTGGAGCAGGGTGGTCACACTGACCTTGCAGAGTTCTGAGAATCAGAGACAGAACATA** 14725400
14725401 **AAAGGCCTGGAAAACATTCTCCAAAAAGAAGCTGCAACATGTGTGGACAATGGGCTTTTC** 14725460
14725461 **ATGCCTCTCTTACTGTCTCTTACTGTC**TATTGACCTGGTGCAAGAAACATGCTCTGGTGA 14725520
14725521 TGGCTGTGAGGGAGGAATGAGGATAGACATAGACACTCCTGTGTCTCAAACATGCTTCTT 14725580
14725581 TATTACTCTGTTATGACTCTGTCTTCCCTGGGGCAGGACCCCAGCCTGCCTACATTTGCA 14725640
14725641 GACAGACACAGTGGCATGTGGAGACAACAGTGTGTCCCAATGACTTTTCTTTACCCCCCA 14725700
14725701 GCTGTCGGCAGTACTCAGTGGAAGGGTGATATTATGACACTGACACTGCTATTTTGAAAC 14725760
14725761 CTGGAGAATGGAAAGGTGCAAAAATCT 14725787

**>chromosome:GRCh38:16:14764161:14764907:1**
14764161 CCTGGGAGTGAAAAGAAATATTACAGCCATGCCTAACTGACTTCTTGAGGTAAGATTGTT 14764220
14764221 CTGTCAGAAAACCCTCTCCCAGTTCCCCTGCAGCTCTTCAGGAATCCACATCTCTCCAGA 14764280
14764281 GCTCTTTGTTCTCATGGGTGGCACCTCCAGAGTGAAGAAGATCCTTTGTCAAGAAGGGAA 14764340
14764341 ACAGAGGGGAAATGAGAGGGTCCTGCAGGCAGAGCTGGAATCAACTTCCACTCTGCCTCT 14764400
14764401 TGCAAGCTGTGTGACCCTGGGCACAATTTCTCCTTCCTCTGGAAACCTCTGTTTTCTTAG 14764460
14764461 **ATTTGGAGCAGGGTGGTCACACTGACCTTGCAGAGTTCTGAGAATCAGAGACAGAACATA** 14764520
14764521 **AAAGGCCTGGAAAACATTCTCCAAAAAGAAGCTGCAACATGTGTGGACAATGGGCTTTTC** 14764580
14764581 **ATGCCTCTCTTACTGTCTCTTACTGTC**TATTGACCTGGTGCAAGAAACATGCTCTGGTGA 14764640
14764641 TGGCTGTGAGGGAGGAATGAGGATAGACATAGACACTCCTGTGTCTCAAACATGCTTCTT 14764700
14764701 TATTACTCTGTTATGACTCTGTCTTCCCTGGGGCAGGACCCCAGCCTGCCTACATTTGCA 14764760
14764761 GACAGACACAGTGGCATGTGGAGACAACAGTGTGTCCCAATGACTTTTCTTTACCCCCCA 14764820
14764821 GCTGTCGGCAGTACTCAGTGGAAGGGTGATATTATGACACTGACACTGCTATTTTGAAAC 14764880
14764881 CTGGAGGATGGAAAGGTGCAAAAATCT 14764907

**>chromosome:GRCh38:16:14950773:14951519:1**
14950773 CCTGGGAGTGAAAAGAAATATTACAGCCATGCCTAACTGACTTCTTGAGGTAAGATTGTT 14950832
14950833 CTGTCAGAAAACCCTCTCCCAGTTCCCCTGCAGCTCTTCAGGAATCCACATCTCTCCAGA 14950892

14950893 GCTCTTTGTTCTCATGGGTGGCACCTCCAGAGTGAAGAAGATCCTTTGTCAAGAAGGGAA 14950952
14950953 ACAGAGGGGAAATGAGAGGGTCCTGCAGGCAGAGCTGGAATCAACTTCCACTCTGCCTCT 14951012
14951013 TGCAAGCTGTGTGACCCTGGGCACAATTTCTCCTTCCTCTGGAAACCTCTGTTTTCTTAG 14951072
14951073 **ATTTGGAGCAGGGTGGTCACACTGACCTTGCAGAGTTCTGAGAATCAGAGACAGAACATA** 14951132
14951133 **AAAGGCCTGGAAAACATTCTCCAAAAAGAAGCTGCAACATGTGTGGACAATGGGCTTTTC** 14951192
14951193 **ATGCCTCTCTTACTGTCTCTTACTGTC**TATTGACCTGGTGCAAGAAACATGCTCTGGTGA 14951252
14951253 TGGCTGTGAGGGAGGAATGAGGATAGACATAGACACTCCTGTGTCTCAAACATGCTTCTT 14951312
14951313 TATTACTCTGTTATGACTCTGTCTTCCCTGGGGCAGGACCCCAGCCTGCCTACATTTGCA 14951372
14951373 GACAGACACAGTGGCATGTGGAGACAACAGTGTGTCCCAATGACTTTTCTTTACCCCCTA 14951432
14951433 GCTGTCGGCAGTACTCAGTGGAAGGGTGATATTATGACACTGACACTGCTATTTTGAAAC 14951492
14951493 CTGGAGGATGGAAAGGTGCAAAAATCT 14951519

**>chromosome:GRCh38:16:16349250:16349996:1**
16349250 CCTGGGAGTGAAAAGAAATATTACAGCCATGCCTAACTGACTTCTTGAGGTAAGATTGTT 16349309
16349310 CTGTCAGAAAACCCTCTCCCAGTTCCCCTGCAGCTCTTCAGGAATCCACATCTCTCCAGA 16349369
16349370 GCTCTTTGTTCTCATGGGTGGCACCTCCAGAGTGAAGAAGATCCTTTGTCAAGAAGGGAA 16349429
16349430 ACAGAGGGGAAATGAGAGGGTCCTGCAGGCAGAGCTGGAATCAACTTCCACTCTGCCTCT 16349489
16349490 TGCAAGCTGTGTGACCCTGGGCACAATTTCTCCTTCCTCTGGAAACCTCTGTTTTCTTAG 16349549
16349550 **ATTTGGAGCAGGGTGGTCACACTGACCTTGCAGAGTTCTGAGAATCAGAGACAGAACATA** 16349609
16349610 **AAAGGCCTGGAAAACATTCTCCAAAAAGAAGCTGCAACATGTGTGGACAATGGGCTTTTC** 16349669
16349670 **ATGCCTCTCTTACTGTCTCTTACTGTC**TATTGACCTGGTGCAAGAAACATGCTCTGGTGA 16349729
16349730 TGGCTGTGAGGGAGGAATGAGGATAGACATAGACACTCCTGTGTCTCAAACATGCTTCTT 16349789
16349790 TATTACTCTGTTATGACTCTGTCTTCCCTGGGGCAGGACCCCAGCCTGCCTACATTTGCA 16349849
16349850 GACAGACACAGTGGCATGTGGAGACAACAGTGTGTCCCAATGACTTTTCTTTACCCCCTA 16349909
16349910 GCTGTCGGCAGTACTCAGTGGAAGGGTGATATTATGACACTGACACTGCTATTTTGAAAC 16349969
16349970 CTGGAGGATGGAAAGGTGCAAAAATCT 16349996

**>chromosome:GRCh38:16:16392614:16393360:1**
16392614 CCTGGGAGTGAAAAGAAATATTACAGCCATGCCTAACTGACTTCTTGAGGTAAGATTGTT 16392673
16392674 CTGTCAGAAAACCCTCTCCCAGTTCCCCTGCAGCTCTTCAGGAATCCACATCTCTCCAGA 16392733
16392734 GCTCTTTGTTCTCATGGGTGGCACCTCCAGAGTGAAGAAGATCCTTTGTCAAGAAGGGAA 16392793
16392794 ACAGAGGGGAAATGAGAGGGTCCTGCAGGCAGAGCTGGAATCAACTTCCACTCTGCCTCT 16392853
16392854 TGCAAGCTGTGTGACCCTGGGCACAATTTCTCCTTCCTCTGGAAACCTCTGTTTTCTTAG 16392913
16392914 **ATTTGGAGCAGGGTGGTCACACTGACCTTGCAGAGTTCTGAGAATCAGAGACAGAACATA** 16392973
16392974 **AAAGGCCTGGAAAACATTCTCCAAAAAGAAGCTGCAACATGTGTGGACAATGGGCTTTTC** 16393033
16393034 **ATGCCTCTCTTACTGTCTCTTACTGTC**TATTGACCTGGTGCAAGAAACATGCTCTGGTGA 16393093
16393094 TGGCTGTGAGGGAGGAATGAGGATAGACATAGACACTCCTGTGTCTCAAACATGCTTCTT 16393153
16393154 TATTACTCTGTTATGACTCTGTCTTCCCTGGGGCAGGACCCCAGCCTGCCTACATTTGCA 16393213
16393214 GACAGACACAGTGGCATGTGGAGACAACAGTGTGTCCCAATGACTTTTCTTTACCCCCTA 16393273
16393274 GCTGTCGGCAGTACTCAGTGGAAGGGTGATATTATGACACTGACACTGCTATTTTGAAAC 16393333
16393334 CTGGAGGATGGAAAGGTGCAAAAATCT 16393360

**>chromosome:GRCh38:16:18358684:18359430:-1**
18359430 CCTGGGAGTGAAAAGAAATATTACAGCCATGCCTAACTGACTTCTTGAGGTAAGATTGTT 18359371
18359370 CTGTCAGAAAACCCTCTCCCAGTTCCCCTGCAGCTCTTCAGGAATCCACATCTCTCCAGA 18359311
18359310 GCTCTTTGTTCTCATGGGTGGCACCTCCAGAGTGAAGAAGATCCTTTGTCAAGAAGGGAA 18359251
18359250 ACAGAGGGGAAATGAGAGGGTCCTGCAGGCAGAGCTGGAATCAACTTCCACTCTGCCTCT 18359191
18359190 TGCAAGCTGTGTGACCCTGGGCACAATTTCTCCTTCCTCTGGAAACCTCTGTTTTCTTAG 18359131
18359130 **ATTTGGAGCAGGGTGGTCACACTGACCTTGCAGAGTTCTGAGAATCAGAGACAGAACATA** 18359071
18359070 **AAAGGCCTGGAAAACATTCTCCAAAAAGAAGCTGCAACATGTGTGGACAATGGGCTTTTC** 18359011
18359010 **ATGCCTCTCTTACTGTCTCTTACTGTC**TATTGACCTGGTGCAAGAAACATGCTCTGGTGA 18358951
18358950 TGGCTGTGAGGGAGGAATGAGGATAGACATAGACACTCCTGTGTCTCAAACATGCTTCTT 18358891
18358890 TATTACTCTGTTATGACTCTGTCTTCCCTGGGGCAGGACCCCAGCCTGCCTACATTTGCA 18358831
18358830 GACAGACACAGTGGCATGTGGAGACAACAGTGTGTCCCAATGACTTTTCTTTACCCCCTA 18358771
18358770 GCTGTCGGCAGTACTCAGTGGAAGGGTGATATTATGACACTGACACTGCTATTTTGAAAC 18358711
18358710 CTGGAGGATGGAAAGGTGCAAAAATCT 18358684

**>chromosome:GRCh38:16:18318531:18319277:-1**
18319277 CCTGGGAGTGAAAAGAAATATTACAGCCATGCCTAACTGACTTCTTGAGGTAAGATTGTT 18319218
18319217 CTGTCAGAAAACCCTCTCCCAGTTCCCCTGCAGCTCTTCAGGAATCCACATCTCTCCAGA 18319158
18319157 GCTCTTTGTTCTCATGGGTGGCACCTCCAGAGTGAAGAAGATCCTTTGTCAAGAAGGGAA 18319098
18319097 ACAGAGGGGAAATGAGAGGGTCCTGCAGGCAGAGCTGGAATCAACTTCCACTCTGCCTCT 18319038
18319037 TGCAAGCTGTGTGACCCTGGGCACAATTTCTCCTTCCTCTGGAAACCTCTGTTTTCTTAG 18318978
18318977 **ATTTGGAGCAGGGTGGTCACACTGACCTTGCAGAGTTCTGAGAATCAGAGACAGAACATA** 18318918

```
18318917 AAAGGCCTGGAAAACATTCTCCAAAAAGAAGCTGCAACATGTGTGGACAATGGGCTTTTC 18318858
18318857 ATGCCTCTCTTACTGTCTCTTACTGTCTATTGACCTGGTGCAAGAAACATGCTCTGGTGA 18318798
18318797 TGGCTGTGAGGGAGGAATGAGGATAGACATAGACACTCCTGTGTCTCAAACATGCTTCTT 18318738
18318737 TATTACTCTGTTATGACTCTGTCTTCCCTGGGGCAGGACCCCAGCCTGCCTACATTTGCA 18318678
18318677 GACAGACACAGTGGCATGTGGAGACAACAGTGTGTCCCAATGACTTTTCTTTACCCCCTA 18318618
18318617 GCTGTCGGCAGTACTCAGTGGAAGGGTGATATTATGACACTGACACTGCTATTTTGAAAC 18318558
18318557 CTGGAGGATGGAAAGGTGCAAAAATCT                                  18318531
```

**>chromosome:GRCh38:16:15364165:15364911:-1**

```
15364911 CCTGGGAGTGAAAAGAAATATTACAGCCATGCCTAACTGACTTCTTGAGGTAAGATTGTT 15364852
15364851 CTGTCAGAAAACCCTCTCCCAGTTCCCCTGCAGCTCTTCAGGAATCCACATCTCTCCAGA 15364792
15364791 GCTCTTTGTTCTCATGGGTGGCACCTCCAGAGTGAAGAAGATCCTTTGTCAAGAAGGGAA 15364732
15364731 ACAGAGGGGAAATGAGAGGGTCCTGCAGGCAGAGCTGGAATCAACTTCCACTCTGCCTCT 15364672
15364671 TGCAAGCTGTGTGACCCTGGGCACAATTTCTCCTTCCTCTGGAAACCTCTGTTTTCTTAG 15364612
15364611 ATTTGGAGCAGGGTGGTCACACTGACCTTGCAGAGTTCTGAGAATCAGAGACAGAACATA 15364552
15364551 AAAGGCCTGGAAAACATTCTCCAAAAAGAAGCTGCAACATGTGTGGACAATGGGCTTTTC 15364492
15364491 ATGCCTCTCTTACTGTCTCTTACTGTCTATTGACCTGGTGCAAGAAACATGCTCTGGTGA 15364432
15364431 TGGCTGTGAGGGAGGAATGAGGATAGACATAGACACTCCTGTGTCTCAAACATGCTTCTT 15364372
15364371 TATTACTCTGTTATGACTCTGTCTTCCCTGGGGCAGGACCCCAGCCTGCCTACATTTGCA 15364312
15364311 GACAGACACAGTGGCATGTGGAGACAACAGTGTGTCCCAATGACTTTTCTTTACCCCCCA 15364252
15364251 GTTGTCGGCAGTACTCAGTGGAAGGGTGATATTATGACACTGACACTGCTATTTTGAAAC 15364192
15364191 CTGGAGGATGGAAAGGTGCAAAAATCT                                  15364165
```

**>scaffold:GRCh38:HSCHR16_1_CTG1:297904:298650:1**

```
297904 CCTGGGAGTGAAAAGAAATATTACAGCCATGCCTAACTGACTTCTTGAGGTAAGATTGTT 297963
297964 CTGTCAGAAAACCCTCTCCCAGTTCCCCTGCAGCTCTTCAGGAATCCACATCTCTCCAGA 298023
298024 GCTCTTTGTTCTCATGGGTGGCACCTCCAGAGTGAAGAAGATCCTTTGTCAAGAAGGGAA 298083
298084 ACAGAGGGGAAATGAGAGGGTCCTGCAGGCAGAGCTGGAATCAACTTCCACTCTGCCTCT 298143
298144 TGCAAGCTGTGTGACCCTGGGCACAATTTCTCCTTCCTCTGGAAACCTCTGTTTTCTTAG 298203
298204 ATTTGGAGCAGGGTGGTCACACTGACCTTGCAGAGTTCTGAGAATCAGAGACAGAACATA 298263
298264 AAAGGCCTGGAAAACATTCTCCAAAAAGAAGCTGCAACATGTGTGGACAATGGGCTTTTC 298323
298324 ATGCCTCTCTTACTGTCTCTTACTGTCTATTGACCTGGTGCAAGAAACATGCTCTGGTGA 298383
298384 TGGCTGTGAGGGAGGAATGAGGATAGACATAGACACTCCTGTGTCTCAAACATGCTTCTT 298443
298444 TATTACTCTGTTATGACTCTGTCTTCCCTGGGGCAGGACCCCAGCCTGCCTACATTTGCA 298503
298504 GACAGACACAGTGGCATGTGGAGACAACAGTGTGTCCCAATGACTTTTCTTTACCCCCCA 298563
298564 GCTGTCGGCAGTACTCAGTGGAAGGGTGATATTATGACACTGACACTGCTATTTTGAAAC 298623
298624 CTGGAGGATGGAAAGGTGCAAAAATCT                                  298650
```

**>scaffold:GRCh38:HSCHR16_1_CTG1:564146:564892:1**

```
564146 CCTGGGAGTGAAAAGAAATATTACAGCCATGCCTAACTGACTTCTTGAGGTAAGATTGTT 564205
564206 CTGTCAGAAAACCCTCTCCCAGTTCCCCTGCAGCTCTTCAGGAATCCACATCTCTCCAGA 564265
564266 GCTCTTTGTTCTCATGGGTGGCACCTCCAGAGTGAAGAAGATCCTTTGTCAAGAAGGGAA 564325
564326 ACAGAGGGGAAATGAGAGGGTCCTGCAGGCAGAGCTGGAATCAACTTCCACTCTGCCTCT 564385
564386 TGCAAGCTGTGTGACCCTGGGCACAATTTCTCCTTCCTCTGGAAACCTCTGTTTTCTTAG 564445
564446 ATTTGGAGCAGGGTGGTCACACTGACCTTGCAGAGTTCTGAGAATCAGAGACAGAACATA 564505
564506 AAGGGCCTGGAAAACATTCTCCAAAAAGAAGCTGCAACATGTGTGGACAATGGGCTTTTC 564565
564566 ATGCCTCTCTTACTGTCTCTTACTGTCTATTGACCTGGTGCAAGAAACATGCTCTGGTGA 564625
564626 TGGCTGTGAGGGAGGAATGAGGATAGACATAGACACTCCTGTGTCTCAAACATGCTTCTT 564685
564686 TATTACTCTGTTATGACTCTGTCTTCCCTGGGGCAGGACCCCAGCCTGCCTACATTTGCA 564745
564746 GACAGACACAGTGGCATGTGGAGACAACAGTGTGTCCCAGTGACTTTTCTTTACCCCCCA 564805
564806 GCTGTCGGCAGTACTCAGTGGAAGGGTGATATGACACTGATACTGCTATTTTGAAACCTG 564865
564866 GAGGATGGAAAGGTGCAAAAATCTATC                                  564892
```

**>scaffold:GRCh38:HSCHR16_1_CTG1:1022111:1022857:-1**

```
1022857 CCTGGGAGTGAAAAGAAATATTACAGCCATGCCTAACTGACTTCTTGAGGTAAGATTGTT 1022798
1022797 CTGTCAGAAAACCCTCTCCCAGTTCCCCTGCAGCTCTTCAGGAATCCACATCTCTCCAGA 1022738
1022737 GCTCTTTGTTCTCATGGGTGGCACCTCCAGAGTGAAGAAGATCCTTTGTCAAGAAGGGAA 1022678
1022677 ACAGAGGGGAAATGAGAGGGTCCTGCAGGCAGAGCTGGAATCAACTTCCACTCTGCCTCT 1022618
1022617 TGCAAGCTGTGTGACCCTGGGCACAATTTCTCCTTCCTCTGGAAACCTCTGTTTTCTTAG 1022558
1022557 ATTTGGAGCAGGGTGGTCACACTGACCTTGCAGAGTTCTGAGAATCAGAGACAGAACATA 1022498
1022497 AAAGGCCTGGAAAACATTCTCCAAAAAGAAGCTGCAACATGTGTGGACAATGGGCTTTTC 1022438
1022437 ATGCCTCTCTTACTGTCTCTTACTGTCTATTGACCTGGTGCAAGAAACATGCTCTGGTGA 1022378
1022377 TGGCTGTGAGGGAGGAATGAGGATAGACATAGACACTCCTGTGTCTCAAACATGCTTCTT 1022318
1022317 TATTACTCTGTTATGACTCTGTCTTCCCTGGGGCAGGACCCCAGCCTGCCTACATTTGCA 1022258
```

1022257 GACAGACACAGTGGCATGTGGAGACAACAGTGTGTCCCAATGACTTTTCTTTACCCCCCA 1022198
1022197 GCTGTCGGCAGTACTCAGTGGAAGGGTGATATTATGACACTGACACTGCTATTTTGAAAC 1022138
1022137 CTGGAGGATGGAAAGGTGCAAAAATCT 1022111

**>scaffold:GRCh38:HSCHR16_1_CTG1:2008440:2009186:1**
2008440 CCTGGGAGTGAAAAGAAATATTACAGCCATGCCTAACTGACTTCTTGAGGTAAGATTGTT 2008499
2008500 CTGTCAGAAAACCCTCTCCCAGTTCCCCTGCAGCTCTTCAGGAATCCACATCTCTCCAGA 2008559
2008560 GCTCTTTGTTCTCATGGGTGGCACCTCCAGAGTGAAGAAGATCCTTTGTCAAGAAGGGAA 2008619
2008620 ACAGAGGGGAAATGAGAGGGTCCTGCAGGCAGAGCTGGAATCAACTTCCACTCTGCCTCT 2008679
2008680 TGCAAGCTGTGTGACCCTGGGCACAATTTCTCCTTCCTCTGGAAACCTCTGTTTTCTTAG 2008739
2008740 **ATTTGGAGCAGGGTGGTCACACTGACCTTGCAGAGTTCTGAGAATCAGAGACAGAACATA** 2008799
2008800 **AAAGGCCTGGAAAACATTCTCCAAAAAGAAGCTGCAACATGTGTGGACAATGGGCTTTTC** 2008859
2008860 **ATGCCTCTCTTACTGTCTCTTACTGTC**TATTGACCTGGTGCAAGAAACATGCTCTGGTGA 2008919
2008920 TGGCTGTGAGGGAGGAATGAGGATAGACATAGACACTCCTGTGTCTCAAACATGCTTCTT 2008979
2008980 TATTACTCTGTTATGACTCTGTCTTCCCTGGGGCAGGACCCCAGCCTGCCTACATTTGCA 2009039
2009040 GACAGACACAGTGGCATGTGGAGACAACAGTGTGTCCCAATGACTTTTCTTTACCCCCTA 2009099
2009100 GCTGTCGGCAGTACTCAGTGGAAGGGTGATATTATGACACTGACACTGCTATTTTGAAAC 2009159
2009160 CTGGAGGATGGAAAGGTGCAAAAATCT 2009186

**>scaffold:GRCh38:HSCHR16_1_CTG1:2053656:2054402:1**
2053656 CCTGGGAGTGAAAAGAAATATTACAGCCATGCCTAACTGACTTCTTGAGGTAAGATTGTT 2053715
2053716 CTGTCAGAAAACCCTCTCCCAGTTCCCCTGCAGCTCTTCAGGAATCCACATCTCTCCAGA 2053775
2053776 GCTCTTTGTTCTCATGGGTGGCACCTCCAGAGTGAAGAAGATCCTTTGTCAAGAAGGGAA 2053835
2053836 ACAGAGGGGAAATGAGAGGGTCCTGCAGGCAGAGCTGGAATCAACTTCCACTCTGCCTCT 2053895
2053896 TGCAAGCTGTGTGACCCTGGGCACAATTTCTCCTTCCTCTGGAAACCTCTGTTTTCTTAG 2053955
2053956 **ATTTGGAGCAGGGTGGTCACACTGACCTTGCAGAGTTCTGAGAATCAGAGACAGAACATA** 2054015
2054016 **AAAGGCCTGGAAAACATTCTCCAAAAAGAAGCTGCAACATGTGTGGACAATGGGCTTTTC** 2054075
2054076 **ATGCCTCTCTTACTGTCTCTTACTGTC**TATTGACCTGGTGCAAGAAACATGCTCTGGTGA 2054135
2054136 TGGCTGTGAGGGAGGAATGAGGATAGACATAGACACTCCTGTGTCTCAAACATGCTTCTT 2054195
2054196 TATTACTCTGTTATGACTCTGTCTTCCCTGGGGCAGGACCCCAGCCTGCCTACATTTGCA 2054255
2054256 GACAGACACAGTGGCATGTGGAGACAACAGTGTGTCCCAATGACTTTTCTTTACCCCCTA 2054315
2054316 GCTGTCGGCAGTACTCAGTGGAAGGGTGATATTATGACACTGACACTGCTATTTTGAAAC 2054375
2054376 CTGGAGGATGGAAAGGTGCAAAAATCT 2054402

**>chromosome:GRCh38:16:28657186:28657935:1**
28657186 CCTGGGAGTGAAAAGAAATATTACAGCCATGCCTAAGTGACTTCTTGAGGTGAGATTGTT 28657245
28657246 CTGTCAGAAAACCCTCTCCCAGTTCCCCTGCAGCTCTTCAGGAATCCACATCTCTCCAGA 28657305
28657306 GCTCTTTGTTCTCATGGGTGGCACCTCCAGAGTGAAGAAGTTCCTTTGTCAAGAAGGGAA 28657365
28657366 ACAGAGGGGAAATGAGAGGGTCCTGCAGGCAGAGCTGGAATCAACTTCCACTCTGCCTCT 28657425
28657426 TGCAAGCTGTGTGACCCTGGGCACAATTTCTCCTTCCTCTGGAAACCTCTGTTTTCTTAG 28657485
28657486 **ATTTGGAGCAGGGTGGTCACACTGACCTTGCAGAGTTCTGAGAGTCAGAGACAGAACATG** 28657545
28657546 **AAAGGCCTGGAAAACATTCTCCAAAAAGAAGCTGCAACATGTGTGGACAATGGGCTTTTC** 28657605
28657606 **ATGCCTCTCTTACTGTCTCTTACTGTCTGT**TGACCTGGTGCAAGAAACATGCTCTGGTGA 28657665
28657666 TGGCTGTGAGGGAGGAATGAGGATAGACATAGACACTCCTGTGTCTCAAACATGCTTCTT 28657725
28657726 TATTACTCTGTTATGACTCTGTCTTCCCTGGGGCAGGACCCCAGCCTGCCTACATTTGCA 28657785
28657786 GACAGACACAGTGGCATGTGGAGACAACAGTGTGTCCCAATGACTTTCCTTTACCCTCCA 28657845
28657846 GCTGTCGGCAGTACTCAGTGGAAGGGTGATATTATGACACTGATACTGCTATTTTGAAAC 28657905
28657906 CTGGAGGATGGAAAGGTGCAAAAATCTATC 28657935

**>chromosome:GRCh38:16:28457119:28457868:-1**
28457868 CCTGGGAGTGAAAAGAAATATTACAGCCATGCCTAAGTGACTTCTTGAGGTGAGATTGTT 28457809
28457808 CTGTCAGAAAACCCTCTCCCAGTTCCCCTGCAGCTCTTCAGGAATCCACATCTCTCCAGA 28457749
28457748 GCTCTTTGTTCTCATGGGTGGCACCTCCAGAGTGAAGAAGATCCTTTGTCAAGAAGGGAA 28457689
28457688 ACAGAGGGGAAATGAGAGGGTCCTGCAGGCAGAGCTGGAATCAACTTCCACTCTGCCTCT 28457629
28457628 TGCAAGCTGTGTGACCCTGGGCACAATTTCTCCTTCCTCTGGAAACCTCTGTTTTCTTAG 28457569
28457568 **ATTTGGAGCAGGGTGGTCACACTGACCTTGCAGAGTTCTGAGAGTCAGAGACAGAACATG** 28457509
28457508 **AAAGGCCTGGAAAACATTCTCCAAAAAGAAGCTGCAACATGTGTGGACAATGGGCTTTTC** 28457449
28457448 **ATGCCTCTCTTACTGTCTCTTACTGTCTGT**TGACCTGGTGCAAGAAACATGCTCTGGTGA 28457389
28457388 TGGCTGTGAGGGAGGAATGAGGATAGACATAGACACTCCTGTGTCTCAAACATGCTTCTT 28457329
28457328 TATTACTCTGTTATGACTCTGTCTTCCCTGGGGCAGGACCCCAGCCTGCCTACATTTGCA 28457269
28457268 GACAGACACAGTGGCATGTGGAGACAACAGTGTGTCCCAATGACTTTTCTTTACCCCCCA 28457209
28457208 GCTGTCGGCAGTACTCAGTGGAAGGGTGATATTATGACACTGATACTGCTATTTTGAAAC 28457149
28457148 CTGGAGGATGGAAAGGTGCAAAAATCTATC 28457119

**>chromosome:GRCh38:16:30225896:30226645:-1**
30226645 CCTGGGAGTGAAAAGAAATATTACAGCCGTGCCTAAGTGACTTCTTGAGGTGAGATTGTT 30226586
30226585 CTGTCAGAAAACCCTCTCCCAGTTCCCCTGCAGCTCTTCAGGAATCCACATCTCTCCAGA 30226526
30226525 GCTCTTTGTTCTCATGGGTGGCACCTCCAGAGTGAAGAAGATCCTTTGTCAAGAAGGGAA 30226466
30226465 ACAGAGGGGAAATGAGAGGGTCCTGCAGGCAGAGCTGGAATCAACTTCCACTCTGCCTCT 30226406
30226405 TGCAAGCTGTGTGACCCTGGGCACAATTTCTCCTTCCTCTGGAAACCTCTGTTTTCTTAG 30226346
30226345 **ATTTGGAGCAGGGTGGTCACACTGACCTTGCAGAGTTCTGAGAGTCAGAGACAGAACATA** 30226286
30226285 **AAAGGCCTGGAAAACATTCTCCAAAAAGAAGCTGCAACATGTGTGGACAGTGGGCTTTTC** 30226226
30226225 **ATGCCTCTCTTACTGTCTCTTACTGTCTGT**TGACCTGGTGCAAGAAACATGCTCTGGTGA 30226166
30226165 TGGCTGTGAGGGAGGAATGAGGATAGACATAGACACTCCTGTGTCTCAAACATGCTTCTT 30226106
30226105 TATTACTCTGTTATGACTCTGTCTTCCCTGGGGCAGGACCCCAGCCTGCCTACATTTGCA 30226046
30226045 GACAGACACAGTGGCATGTGGAGACAACAGTGTGTCCCAATGACTTTCCTTTACCCTCCA 30225986
30225985 GCTGTCGGCAGTACTCAGTGGAAGGGTGATATTATGACACTGATACTGCTATTTTGAAAC 30225926
30225925 CTGGAGGATGGAAAGGTGCAAAAATCTATC 30225896

**>chromosome:GRCh38:16:29485926:29486675:-1**
29486675 CCTGGGAGTGAAAAGAAATATTACAGCCATGCCTAAGTGACTTCTTGAGGTGAGATTGTT 29486616
29486615 CTGTCAGAAAACCCTCTCCCAGTTCCCCTGCAGCTCTTCAGGAATCCACATCTCTCCAGA 29486556
29486555 GCTCTTTGTTCTCATGGGTGGCACCTCCAGAGTGAAGAAGATCCTTTGTCAAGAAGGGAA 29486496
29486495 ACAGAGGGGAAATGAGAGGGTCCTGCAGGCAGAGCTGGAATCAACTTCCACTCTGCCTCT 29486436
29486435 TGCAAGCTGTGTGACCCTGGGCACAATTTCTCCTTCCTCTGGAAACCTCTGTTTTCTTAG 29486376
29486375 **ATTTGGAGCAGGGTGGTCACACTGACCTTGCAGAGTTCTGAGAGTCAGAGACAGAACATA** 29486316
29486315 **AAAGGCCTGGAAAACATTCTCCAAAAAGAAGCTGCAACATGTGTGGACAGTGGGCTTTTC** 29486256
29486255 **ATGCCTCTCTTACTGTCTCTTACTGTCTGT**TGACCTGGTGCAAGAAACATGCTCTGGTGA 29486196
29486195 TGGCTGTGAGGGAGGAATGAGGATAGACATAGACACTCCTGTGTCTCAAACATGCTTCTT 29486136
29486135 TATTACTCTGTTATGACTCTGTCTTCCCTGGGGCAGGACCCCAGCCTGCCTACATTTGCA 29486076
29486075 GACAGACACAGTGGCATGTGGAGACAACAGTGTGTCCCAATGACTTTCCTTTACCCTCCA 29486016
29486015 GCTGTCGGCAGTACTCAGTGGAAGGGTGATATTATGACACTGATACTGCTATTTTGAAAC 29485956
29485955 CTGGAGGATGGAAAGGTGCAAAAATCTATC 29485926

**>scaffold:GRCh38:HSCHR16_1_CTG1:790104:790850:1**
790104 CCTGGGAGTGAAAAGAAATATTACAGCCATGCCTAACTGACTTCTTGAGGTAAGATTGTT 790163
790164 CTGTCAGAAAACCCTCTCCCAGTTCCCCTGCAGCTCTTCAGGAATCCACATCTCTCCAGA 790223
790224 GCTCTTTGTTCTCATGGGTGGCACCTCCAGAGTGAAGAAGATCCTTTGTCAAGAAGGGAA 790283
790284 ACAGAGGGGAAATGAGAGGGTCCTGCAGGCAGAGCTGGAATCAACTTCCACTCTGCCTCT 790343
790344 TGCAAGCTGTGTGACCCTGGGCACAATTTCTCCTTCCTCTGGAAACCTCTGTTTTCTTAG 790403
790404 **ATTTGGAGCAGGATGGTCACACTGACCTTGCAGAGTTCTGAGAATCAGAGACAGAACATA** 790463
790464 **AAAGGCCTGGAAAACATTCTCCAAAAAGAAGCTGCAACATGTGTGGACAATGGGCTTTTC** 790523
790524 **ATGCCTCTCTTACTGTCTCTTACTGTC**TATTGACCTGGTGCAAGAAACATGCTCTGGTGA 790583
790584 TGGCTGTGAGGGAGGAATGAGGATAGACATAGACACTCCTGTGTCTCAAACATGCTTCTT 790643
790644 TATTACTCTGTTATGACTCTGTCTTCCCTGGGGCAGGACCCCAGCCTGCCTACATTTGCA 790703
790704 GACAGACACAGTGGCATGTGGAGACAACAGTGTGTCCCAAAGACTTTTCTTTACCCCCTA 790763
790764 GCTGTCGGCAGTACTCAGTGGAAGGGTGATATTATGACACTGACACTGCTATTTTGAAAC 790823
790824 CTGGAGGATGGAAAGGTGCAAAAATCT 790850

**>scaffold:GRCh38:HSCHR16_1_CTG1:718213:718959:-1**
718959 CCTGGGAGTGAAAAGAAATATTACAGCCATGCCTAACTGACTTCTTGAGGTAAGATTGTT 718900
718899 CTGTCAGAAAACCCTCTCCCAGTTCCCCTGCAGCTCTTCAGGAATCCACATCTCTCCAGA 718840
718839 GCTCTTTGTTCTCATGGGTGGCACCTCCAGAGTGAAGAAGATCCTTTGTCAAGAAGGGAA 718780
718779 ACAGAGGGGAAATGAGAGGGTCCTGCAGGCAGAGCTGGAATCAACTTCCACTCTGCCTCT 718720
718719 TGCAAGCTGTGTGACCCTGGGCACAATTTCTCCTTCCTCTGGAAACCTCTGTTTTCTTAG 718660
718659 **ATTTGGAGCAGGATGGTCACACTGACCTTGCAGAGTTCTGAGAATCAGAGACAGAACATA** 718600
718599 **AAAGGCCTGGAAAACATTCTCCAAAAAGAAGCTGCAACATGTGTGGACAATGGGCTTTTC** 718540
718539 **ATGCCTCTCTTACTGTCTCTTACTGTC**TATTGACCTGGTGCAAGAAACATGCTCTGGTGA 718480
718479 TGGCTGTGAGGGAGGAATGAGGATAGACATAGACACTCCTGTGTCTCAAACATGCTTCTT 718420
718419 TATTACTCTGTTATGACTCTGTCTTCCCTGGGGCAGGACCCCAGCCTGCCTACATTTGCA 718360
718359 GACAGACACAGTGGCATGTGGAGACAACAGTGTGTCCCAAAGACTTTTCTTTACCCCCTA 718300
718299 GCTGTCGGCAGTACTCAGTGGAAGGGTGATATTATGACACTGACACTGCTATTTTGAAAC 718240
718239 CTGGAGGATGGAAAGGTGCAAAAATCT 718213

**>chromosome:GRCh38:16:69976824:69977564:-1**
69977564 TACCTGGGAGTGAAAAGAAATATTACAGCCATGCCTAACTGACTTCTTGAGGTAAGATTG 69977505
69977504 TTCTGTCAGAAAACCCTCTCCCAGTTCCCCTGCAGCTCTTCAGGAATCCACATCTCTGCA 69977445
69977444 GAGCTCTTTGTTCTCATGGGTGGCACCTCCAGAGTGAAGAAGATCCTTTATCAAGAAGGG 69977385

69977384 AAACAGGGGAAATGAGAGGGTCCTGCAGGCAGAGCTGGAATCAACTTCCACTCTGCCTCT 69977325
69977324 TGCAAGCTGTGTGACCCCGGGCACAATTTCTCCTTCCTCTGGAAACCTCTGTTTTCTTAG 69977265
69977264 **ATTTGGAGCAGGGTGGTCACACTGACCTTGCAGAGTTCCGAGAATCAGAGACAGAACATA** 69977205
69977204 **AAAGGCCTGGAAAACATTCTCCAAAAAGAAGCTGCAACATGTGTGGACAATGGGCTTTTC** 69977145
69977144 **ATGCCTCTCTTACTGTCTGTT**GACCTGGTGCAAGAAACATGCTCTGGTGATGGCTGTGAG 69977085
69977084 GGAGGAATGAGGATAGACATAGACACTCCTGTGTCTCAAACATGCCTCTTTATTACTCTG 69977025
69977024 TTATGACTCTGTCTTCCCTGGGGCAGGACCCCAGCCTGCCTACATTTGCAGACAGACACA 69976965
69976964 GTGGCATGTGGAGACAACAGTGTGTCCCAATGACTTTTCTTTACTCCCCAGCTGTCGGCA 69976905
69976904 GTACTCAGTGGAAGGGTGATATTATGACACTGATACTGCTATTTTGAAACCTGGAGGATG 69976845
69976844 GAAAGGTGCAAAAATCTATCA 69976824

**>chromosome:GRCh38:18:11620012:11620752:-1**
11620752 CCTGGGAGTGAAAAGAAATATTACAGCCATGCCCAACTGACTTCTTGAGGTAAGATTGTT 11620693
11620692 CTGTCAGAAAACCCTCTCCCAGTTCCCCTGCAGCTCTTCAGGAATCCACATCTCTCCAGA 11620633
11620632 GCTCTTTGTTCTCATGGGTGGCACCTCCAGAGTGAAGAAGATCCTTTGTCAAGAAGGGAA 11620573
11620572 ACAGAGGGGAAATGAGAGGGTCCTGCAGGCAGAGCTGGAATCAACTTCCACTCTGCCTCT 11620513
11620512 TGCAAGCTGTGTGACCCTGGGCACAATTTCTCCTTCCTCTGGAAACCTCTGTTTTCTTAG 11620453
11620452 **ATTTGGAGCAGGGTGGTCACACTGACCTTGCAGAGTTCTGAGAATCAGAGACAGAACATA** 11620393
11620392 **AAAGGCCTGGAAAACATTCTCCAAAAAGAAGCTGCAACATGTGTGGACAATGGGCTTTTC** 11620333
11620332 **ATGCCTCTCTTACTGTCTGTT**GACCTGGTGCAAGAAACATGCTCTGGTGATGGCTGTGAG 11620273
11620272 GGAGGAATGAGGATAGACATAGACACTCCTGTGTCTCAAACATGCTTCTTTATTACTCTG 11620213
11620212 CTATGACTCTGTCTTCCCTGGGGCAGGACCCCAGCCTGCCTACATTTGCAGACAGACACA 11620153
11620152 GTGGCATGTGGAGACAACAGTGTGTCCCAATGACTTTTCTTTACCCCCCAGCTGTCGGCA 11620093
11620092 GTACTCAGTGGAAGGGTGATATTATGACACTGATACTGCTATTTTGAAACCTGGAGGATG 11620033
11620032 GAAAGGTGCAAAAATCTATCA 11620012
